# Progenitor-Derived Regulatory Logic Generates Functionally Related Neuronal Subtypes

**DOI:** 10.1101/2024.09.27.615552

**Authors:** Chundi Xu, Rishi G. Sastry, Peter Newstein, Ryan N. Delgado, Neel Wagh, Marion Silies, Constance L. Cepko, Chris Q. Doe

**Author notes:** Co-authors for correspondence at or.

## Abstract

The brain deploys diverse neuronal subtypes to split complex inputs into parallel channels—each tuned to distinct features—enabling rich neural processing. Yet how progenitors generate distinct but functionally related subtypes remains unknown. In the *Drosophila* lamina (five lamina neuron subtypes receiving photoreceptor input), we uncover the regulatory logic: a pan-class homeodomain transcription factor (HDTF), induced by Hedgehog in progenitors and maintained in all lamina neurons, drives diversification within the lamina neuron class by orchestrating a four-step program across the progenitor-to-newborn neuron transition. Specifically, it establishes progenitor identity, promotes cell-cycle exit, induces subtype-specific HDTFs, and acts as their obligate cofactor to specify distinct subtypes. Loss of subtype-specific HDTFs in newborn—but not older—neurons drives subtype-to-subtype fate conversions at molecular, morphological, and functional levels, including a contrast-to-luminance encoding switch. In the mouse retina, we find that each of the 63 amacrine, 15 bipolar, and 45 retinal ganglion cell subtypes expresses pan-class and subtype-specific HDTFs, indicating evolutionary conservation of this regulatory logic. Given the brain-wide expression of HDTFs across species, these findings convert a longstanding mystery into a testable, generalizable principle for within-class subtype diversification and lay the groundwork for subtype-precise reprogramming and cell replacement strategies.

## INTRODUCTION

The brain utilizes many neuronal subtypes to split complex inputs into parallel, specialized channels—each tuned to distinct features—facilitating the simultaneous execution of distinct computations and rich neural processing. Recent single-cell RNA sequencing (scRNA-seq) has systematically mapped the molecular diversity of these subtypes across the brain. For example, in *Drosophila*, 30 olfactory receptor neuron subtypes connect the antenna to the antennal lobe^1^. In the mouse retina, 45 retinal ganglion cell subtypes transform bipolar input into feature-specific signals relayed via the optic nerve to retinorecipient nuclei^2–4^. In the mouse dorsal root ganglia, 14 somatosensory neuron subtypes detect mechanical, thermal, and chemical modalities^5^. In the mouse cerebellum, 11 Purkinje cell subtypes receive input from parallel and climbing fibers and project to the deep cerebellar nuclei^6^. In the mouse midbrain, 20 molecularly distinct dopamine neuron subtypes differ in vulnerability to Parkinson’s disease^7^. Within the mouse hippocampal CA1 region, pyramidal neuron subtypes include those that encode spatial and those that encode non-spatial aspects of the environment^8^. Across these systems, individual neuronal classes comprise multiple molecularly defined subtypes. Yet despite this widespread pattern of within-class subtype diversity, how progenitors generate multiple, functionally related subtypes within a single neuronal class remains unknown, and no general mechanism for such diversification has been established.

We hypothesized that progenitor-derived, pan-class homeodomain transcription factors (HDTFs) induce subtype-specific HDTFs in newborn neurons and then act as their obligate cofactors to specify distinct subtypes within the progenitor-defined class. To test this, we leveraged the genetically tractable *Drosophila* lamina, which contains five lamina neuron subtypes (L1–L5) that process photoreceptor input and transmit signals to the medulla. The developing lamina provides, within one tissue, a spatial continuum from progenitors to newborn and older lamina neurons (Figure 1A), enabling precise dissection of molecular events across the progenitor-to-neuron transition. Using this model, we combined scRNA-seq with stage– and subtype-specific gene perturbations and subtype-specific neural activity assays to define the regulatory logic for within-class subtype diversification, and we tested its conservation in the mouse retina using a single-cell retinal atlas and *in situ* hybridization.

**Figure 1.**
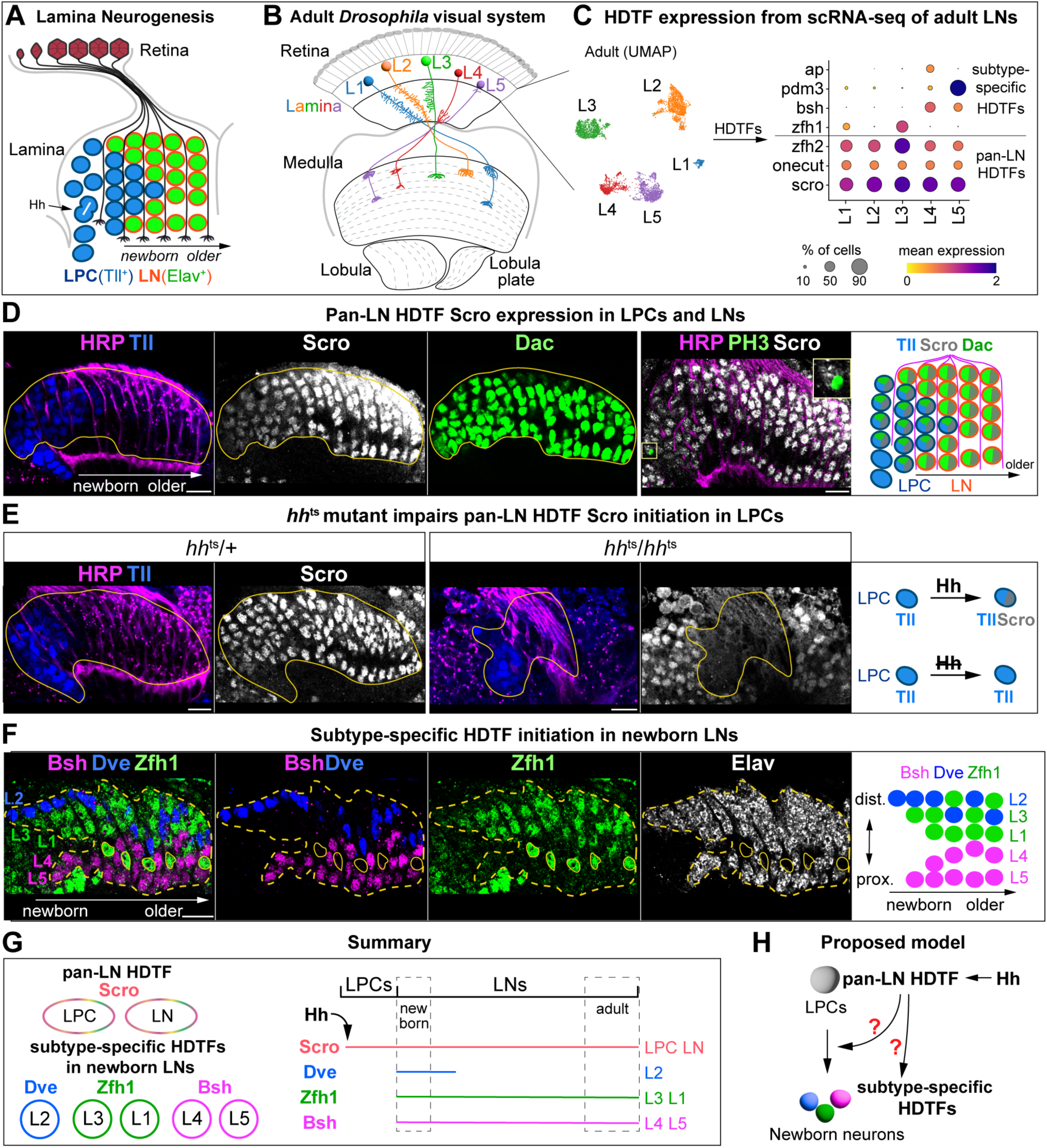
Hedgehog induces the pan-LN HDTF Scro in mitotic LPCs, which is inherited by all newborn lamina neurons, whereas subtype-specific HDTFs (Zfh1, Dve, Bsh) initiate in distinct newborn subsets. (A) Schematic diagram of lamina neurogenesis during later larval and early pupal stages. Photoreceptor axons deliver hedgehog (Hh) to lamina precursor cells (LPCs), triggering their terminal division. Dac is induced by Hh in LPCs and maintained in lamina neurons (LNs). Tailless (Tll) marks LPCs; elav marks LNs. The lamina contains, within a single tissue, a spatial continuum from LPCs to newborn and older LNs. (B) Schematic diagram of the adult *Drosophila* visual system. (C) scRNA-seq of adult LNs identifies two HDTF categories: pan-LN and subtype-specific. UMAP: each point represents one cell; colors denote LN subtypes (Leiden clustering). Dot plot: dot size = percentage of cells in each cluster expressing the gene; color = mean normalized expression in expressing cells. LNs are from 3-day-old adult females. (D) Scro is initiated in mitotic LPCs and maintained in all LNs, showing a pattern similar to Dac. Dac marks a subset of LPCs and all LNs; HRP labels photoreceptor axons. Tll: LPC marker; PH3: mitotic marker. Yellow outlines: Dac^+^ LPCs and LNs; yellow squares: PH3^+^ Scro^+^ mitotic LPC. (E) *hh*^ts^/ *hh*^ts^ mutants fail to initiate Scro expression in LPCs. Yellow outlines: LPCs and LNs. (F) Subtype-specific HDTFs are restricted to distinct newborn LN subsets: Dve in L2; Zfh1 in L1 and L3; Bsh in L4 and L5. Yellow dashed outlines: LNs; yellow circles: Zfh1+ glia. (G) Schematic summary. (H) Proposed model. Scale bar, 10 µm; n ≥ 5 brains; sample age (D-F), 19h after pupal formation (APF). See also Figures S1 and S2.

scRNA-seq, validated by immunostaining, reveals two tiers of HDTFs across the progenitor-to-newborn neuron transition: a pan-class HDTF initiated in progenitors and maintained in all newborn lamina neurons, and subtype-specific HDTFs initiated in newborn lamina neurons. Stage-specific perturbations show that the pan-class HDTF orchestrates a four-step program across this transition: establishing progenitor identity, promoting cell-cycle exit, inducing subtype-specific HDTFs, and acting as their obligate cofactor to specify distinct subtypes. Subtype-specific perturbations, coupled with subtype-specific neural activity assays during parametrically varied luminance and contrast stimuli, reveal that loss of subtype-specific HDTFs in newborn neurons drives subtype-to-subtype fate conversions at molecular, morphological, and functional levels, including a switch from contrast to luminance encoding. In the mouse retina, scRNA-seq validated by *in situ* hybridization reveals pan-bipolar and subtype-specific HDTFs across the 15 bipolar cell subtypes, indicating evolutionary conservation of this regulatory logic. Given the brain-wide expression of HDTFs across species^9–11^, our results define a generalizable regulatory logic for within-class subtype diversification and point to strategies for subtype-precise reprogramming.

## RESULTS

### Two tiers of HDTF expression across the progenitor-to-newborn neuron transition

During late larval and early pupal stages, photoreceptor axons deliver Hedgehog (Hh) to lamina precursor cells (LPCs), inducing expression of the TF Dachshund (Dac) and triggering terminal division^12,13^ (Figure 1A). Following division, postmitotic LPCs align with photoreceptor axons to form columns and differentiate into the five lamina neuron subtypes (L1–L5) within each of the ∼800 cartridges^12^. LPCs are broadly marked by the nuclear receptor Tailless (Tll)^14^, facilitating identification of this population.

To identify HDTFs acting in newborn lamina neurons, we first profiled HDTF expression across adult lamina neurons by single-cell RNA sequencing (scRNA-seq) (Figure 1B and 1C). This analysis yielded two categories: candidate pan-class HDTFs expressed across all five lamina neuron subtypes (pan-LN) and candidate subtype-specific HDTFs restricted to particular subtypes (Figure 1C).

We next asked whether candidate pan-LN HDTFs are expressed during the progenitor-to-newborn neuron transition. Scarecrow (Scro) showed a spatiotemporal pattern resembling Dac: Scro was initiated in mitotic LPCs marked by phospho-Histone H3 (PH3) and persisted in postmitotic LPCs and all lamina neurons (Figure 1D). This suggested that Scro expression may be initiated by Hh signaling. Indeed, blocking Hh activity, using a temperature-sensitive loss-of-function allele (*hh*^ts^), abolished Scro initiation in homozygous mutants (*hh*^ts^/*hh*^ts^), whereas heterozygous controls (*hh*^ts^/+) exhibited normal Scro expression (see Methods) (Figure 1E). The absence of Scro was accompanied by disrupted lamina columns, labeled by horseradish peroxidase (HRP), due to loss of lamina neurogenesis^13^ (Figure 1E). Scro expression persisted from the newborn-neuron stage into adulthood (Figure S1A and S1C). Thus, Scro is a pan-LN HDTF that is induced by Hh in mitotic LPCs and maintained in all lamina neurons. The candidate pan-LN HDTFs Onecut and Zfh2 were not detected during neurogenesis and were excluded from further analysis (Figure S2A and S2B).

We next examined candidate subtype-specific HDTFs during the progenitor-to-newborn neuron transition. Consistent with our previous findings^14^, Zfh1 was expressed in newborn L1 and L3 neurons, and Bsh in newborn L4 and L5 neurons (Figure 1F). By contrast, Ap and Pdm3 were initiated in older L4 and L5 neurons^14^. To identify an L2-specific HDTF, we analyzed a published developing-lamina scRNA-seq dataset^15^ and identified Defective proventriculus (Dve), a previously unreported lamina HDTF. Immunostaining detected Dve protein in newborn L2 and through 2 days after pupal formation (APF), but not at 3 days APF or in adults (Figure 1F, S1B, and S1C). Dve protein was also transiently detected in L5 around 2 days APF but was absent from newborn L5 (Figure 1F, S1B, and S1C). Subtype-specific HDTF expression pattern mapped onto a stereotyped spatial arrangement of newborn lamina neuron somata: the distal layer contained Dve-positive L2 neurons, and intermediate layers contained Zfh1-positive L1/L3 neurons, and proximal layers contained Bsh-positive L4/L5 neurons (Figure 1F). At later stages, L2 and L3 somata became intermingled (Figure 1F). These findings define Zfh1, Dve, and Bsh as subtype-specific HDTFs for newborn L1/L3, L2, and L4/L5, respectively (Figure 1G).

In summary, our data reveal two tiers of HDTF expression across the progenitor-to-newborn neuron transition: (i) a progenitor-derived, pan-LN HDTF (Scro) initiated by Hh in mitotic LPCs and maintained in all newborn lamina neurons, and (ii) newborn-neuron-onset, subtype-specific HDTFs expressed in subtype-restricted patterns (Figure 1G). This temporal ordering supports a model in which Scro promotes LPC cell-cycle exit and initiates subtype-specific HDTF expression in newborn neurons, thereby enabling lamina neuron diversification (Figure 1H).

### Pan-LN HDTF Scro promotes progenitor cell-cycle exit and initiates subtype-specific HDTF expression in newborn neurons

To define the role of the pan-LN HDTF Scro during the progenitor-to-newborn neuron transition, we knocked down Scro (Scro-KD) specifically in LPCs and their lamina neuron progeny (see Methods). Scro-KD nearly abolished Scro expression in both LPCs and lamina neurons (Figure 2A). Knockdown caused a striking expansion of Tll^+^ LPCs before their assembly into HRP-labeled columnar structures, accompanied by a substantial reduction in lamina neurons, indicative of impaired neurogenesis (Figure 2A). Consequently, HRP^+^ lamina columns were significantly smaller, consistent with reduced neuron production and LPC accumulation (Figure 2A). Expanded LPCs retained the mitotic marker phospho-Histone H3 (PH3), indicating that Scro is required for LPC cell-cycle exit (Figure 2A). Consistent with this framework, previously described hypomorphic *scro* mutants lack lamina neurons^16^, and our data provide a mechanistic basis implicating failed LPC cell-cycle exit (Figure 2A).

**Figure 2.**
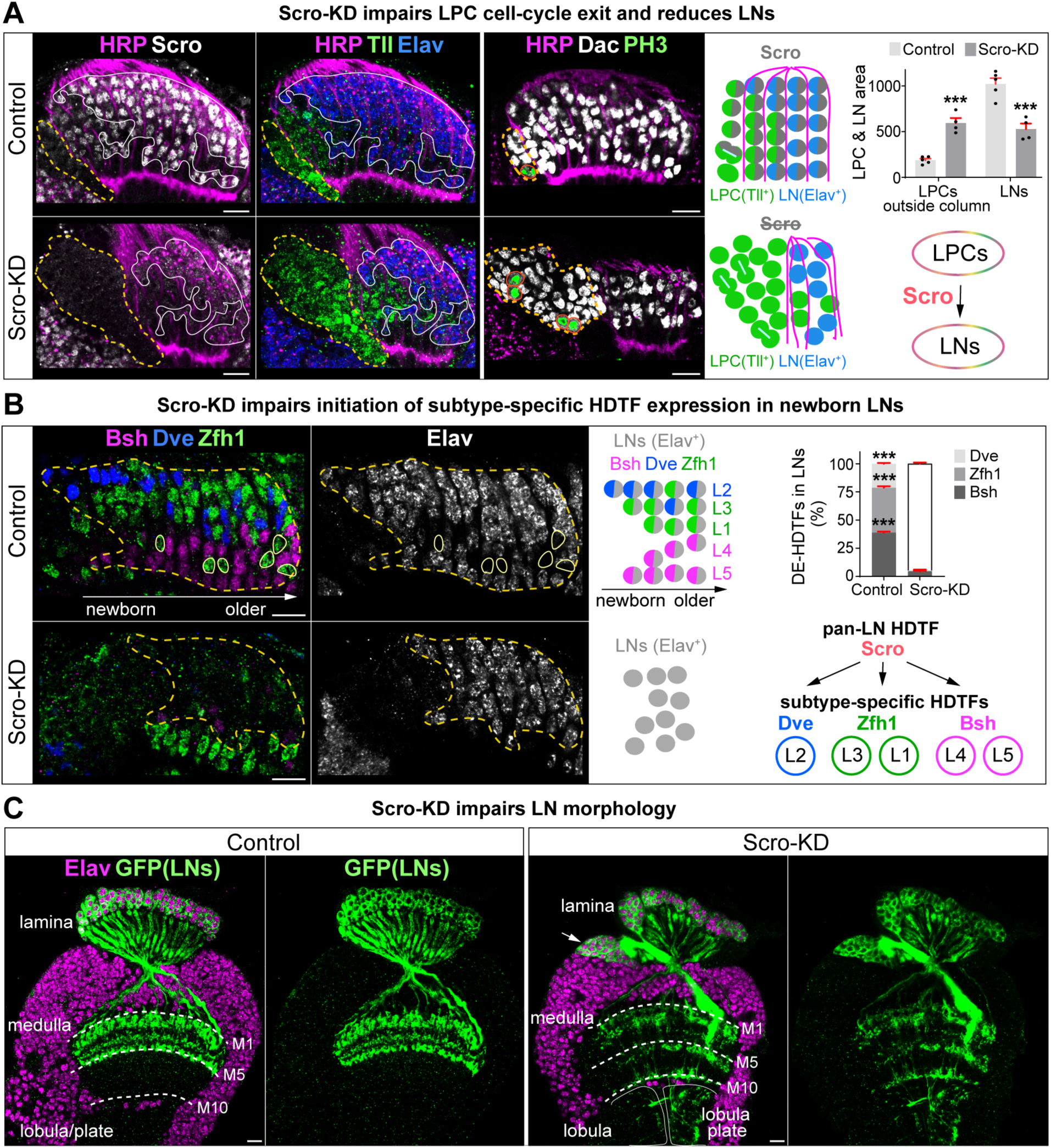
Scro sequentially promotes lamina neurogenesis and initiates expression of Zfh1, Dve, and Bsh in newborn neurons. (A) Scro knockdown (Scro-KD) in LPCs and their neuronal progeny (27G05-Gal4, tub-Gal80^ts^; Scro-RNAi) impairs LPC cell-cycle exit and reduces LNs. Tll: LPC marker; Elav: neuron marker; HRP: lamina columns. Dashed yellow outlines: LPCs outside HRP^+^ columns; white outlines: LNs; orange circles: PH3^+^ mitotic LPCs. Areas of LPCs outside columns and LNs are quantified. Sample age: 19h APF. (B) Scro-KD prevents initiation of Zfh1, Dve, and Bsh in newborn LNs. Yellow dashed outlines: LNs; yellow circles: Zfh1^+^ glia. Percentages of Dve^+^, Zfh1^+^, and Bsh^+^ LNs are quantified. Sample age: 19h APF. (C) Scro-KD disrupts canonical lamina neuron morphology in the mature lamina. Arrow: LN somata fused with medulla neuron somata. N ≥ 5 brains; sample age: 4d APF. Data are mean ± SEM; (A) n = 5 control and n = 4 Scro-KD brains; (B) n = 5 brains per group; \*\*\**p*<0.001, unpaired t-test. Scale bar, 10 µm.

Despite these defects, ∼50% of the normal number of lamina neurons were still generated in Scro-KD animals (Figure 2A), likely due to the delayed onset of knockdown that allowed some LPCs to exit mitosis before substantial Scro depletion. We leveraged this residual neurogenesis to test whether Scro initiates subtype-specific HDTF expression in newborn neurons. In controls, subtype-specific HDTFs were expressed as follows: Zfh1 in L1 and L3, Dve in L2, and Bsh in L4 and L5 (Figure 2B). Scro-KD nearly abolished the expression of all three in newborn lamina neurons, demonstrating that Scro is essential to initiate subtype-specific HDTF expression (Figure 2B).

In the mature lamina, Scro-KD neurons lacking pan-LN and subtype-specific HDTFs exhibited severe morphological defects. Their somata failed to segregate into discrete lamina layers, instead forming aberrant clusters that often merged with medulla neuron somata (Figure 2C). Axonal targeting was profoundly altered: rather than projecting to superficial medulla layers (M1-M5), Scro-KD neurons extended axons broadly across all medulla layers (M1-M10) and into the lobula and lobula plate (Figure 2C). These phenotypes reveal that the pan-LN HDTF Scro is required to establish the class identity of lamina neurons.

Together, these results define a sequential role for Scro during the progenitor-to-newborn neuron transition: establishing progenitor identity, promoting cell-cycle exit, and activating subtype-specific HDTFs in newborn neurons. These data pinpoint the newborn-neuron stage as a critical window in which a progenitor-derived pan-LN HDTF launches the subtype-specific program.

### Subtype-specific HDTFs act in newborn neurons to diversify lamina subtypes

We next asked whether subtype-specific HDTFs act in newborn neurons to drive lamina neuron diversification. As an initial test, we simultaneously knocked down two subtype-specific HDTFs, Bsh and Dve (Bsh/Dve-DKD), reasoning that combined loss might unmask a default neuronal fate. In controls, the three subtype-specific HDTFs showed mutually exclusive expression in newborn lamina neurons: Dve in distal-layer L2 neurons, Bsh in proximal-layer L4 and L5 neurons, and Zfh1 in intermediate-layer L1 and L3 neurons (Figure 3A). In Bsh/Dve-DKD animals, expression of Dve and Bsh was undetectable, and Zfh1 expanded across all newborn lamina neurons, indicating loss of L2, L4, and L5 identities (Figure 3A). As Zfh1 is required for the generation of L1 and L3 neurons^14^, we asked whether ectopic Zfh1 is sufficient to drive these fates. Consistent with Zfh1 expansion, the L1 marker Svp and the L3 marker Erm became broadly expressed across all lamina layers, suggesting subtype conversion from L2, L4, and L5 to either L1 or L3 (Figure 3A). By 4 days APF, the mature lamina in Bsh/Dve-DKD animals comprised only L1 (∼40%) and L3 (∼60%) neurons, confirming persistent fate conversion (Figure S3A). Previously, we showed that Bsh loss alone converts L4 neurons to L3 and L5 neurons to L1^14^. Together, these findings indicate that Bsh and Dve diversify lamina neuron subtypes by repressing default Zfh1-driven L1/L3 fates and promoting alternative fates. They are also consistent with an evolutionary scenario in which an ancestral lamina comprised only L1 and L3 neurons—responsive to increments and decrements of light intensity, respectively^17,18^—with subsequent diversification driven by the acquisition of Bsh and Dve (Figure 3C).

**Figure 3.**
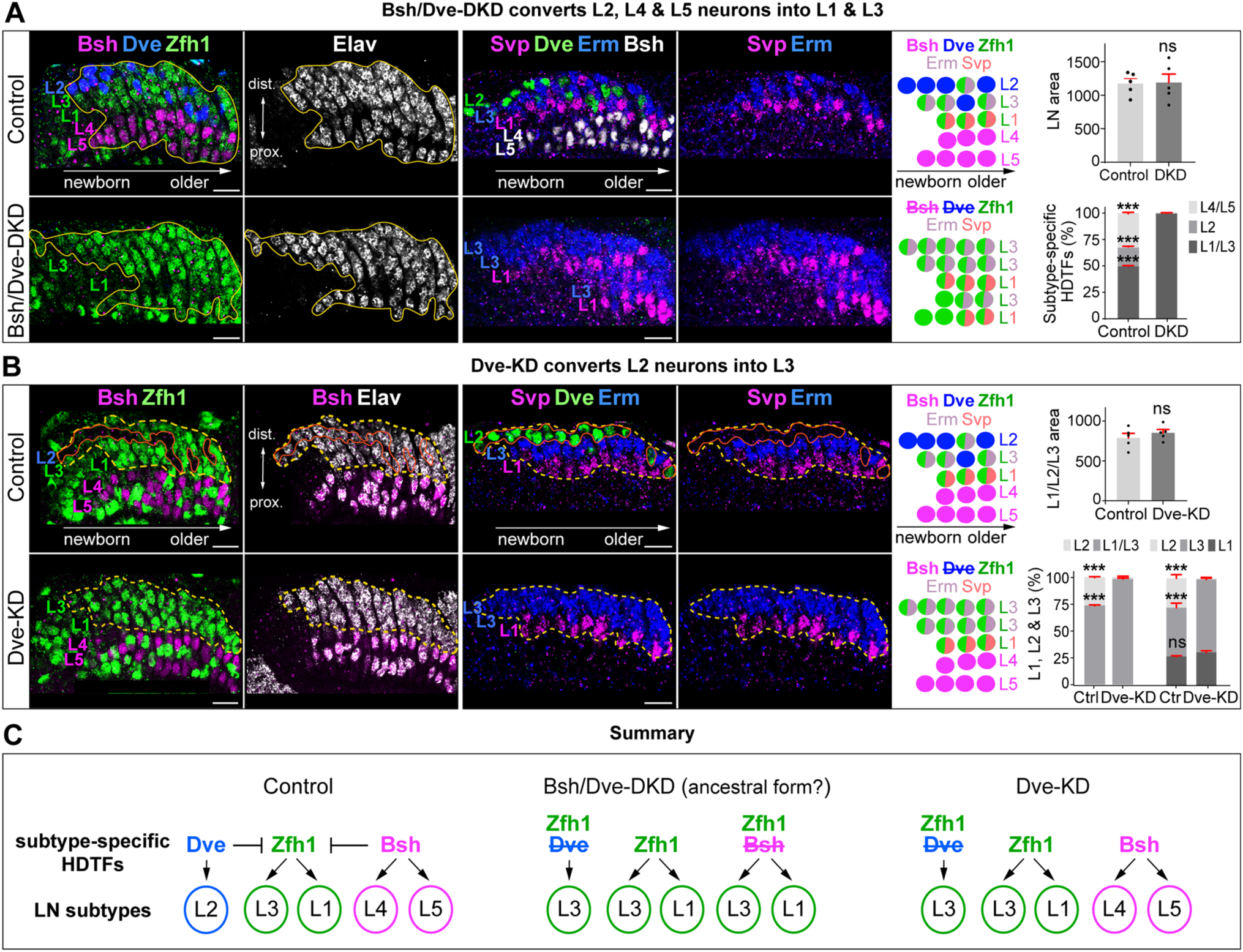
Subtype-specific HDTFs in newborn neurons diversify lamina neuron subtypes. (A) Bsh/Dve double knockdown (Bsh/Dve-DKD) converts L2, L4, and L5 neurons into L1 and L3. DKD was achieved with 27G05-Gal4 driving UAS-Bsh-RNAi and UAS-Dve-dsRNA. Elav: neuron marker; Svp: L1 marker; Erm: L3 marker. Yellow outlines: lamina neurons (LNs). LN area and percentages of L1/L3 (Zfh1^+^Elav^+^), L2 (Dve^+^), and L4/L5 (Bsh^+^) within LN area are quantified. (B) Dve knockdown (Dve-KD) converts L2 neurons into L3. Dve-KD was achieved with 27G05-Gal4 driving UAS-Dve-dsRNA. Yellow dashed outlines: L1, L2, and L3 neurons; orange outlines: L2 neurons. The L1, L2, and L3 neuron area, percentages of L2 (Elav^+^ Zfh1^-^) and L1/L3 (Zfh1^+^) within the L1/L2/L3 area, and percentages of L1 (Svp^+^), L2 (Dve^+^), and L3 (Erm^+^) within the L1/L2/L3 area are quantified. (C) Schematic summary. Our previous work shows that Bsh knockdown induces ectopic Zfh1 and converts L4 and L5 neurons into L3 and L1, respectively^14^. Data are mean ± SEM; n = 5 brains in (A) and (B); ns, not significant; \*\*\**p*<0.001, unpaired t-test. Scale bar, 10 µm; sample age: 19h APF. See also Figures S3 and S4.

To determine the fate adopted by L2 neurons lacking only Dve, we performed lamina-specific Dve knockdown (Dve-KD). Dve-KD abolished Dve expression in L2 neurons and caused Zfh1 to expand into the distal L2 layer, suggesting an L2-to-L1 or L2-to-L3 fate switch (Figure 3B). Subtype marker analysis revealed ectopic Erm (L3 marker), but not Svp (L1 marker), confirming a specific L2-to-L3 conversion (Figure 3B). By 4 days APF, the mature lamina of Dve-KD exhibited an increased proportion of Zfh1^+^/Erm^+^ L3 neurons, accompanied by loss of the L2 marker Bab2, indicating a stable L2-to-L3 conversion (Figure S3B).

Finally, we asked whether subtype-specific HDTFs act specifically at the newborn-neuron stage to specify subtype. The timing of Dve expression provided leverage: Dve is transient in both L2 and L5 neurons, but at different stages—in newborn L2 and in immature L5 (∼2 days APF), and absent from newborn L5 (Figure 1E and S1B). Bsh is expressed in newborn L5, and we previously showed that Bsh-KD converts L5 neurons to L1. Here, L5-specific Dve-KD abolished Dve expression but had no effect on the L5 marker Pdm3, indicating that Dve is dispensable for L5 specification (Figure S4A). Conversely, Pdm3-KD eliminated Dve expression in L5 neurons, placing Pdm3 upstream of Dve and confirming that late Dve expression occurs downstream of subtype specification (Figure S4A and S4B).

Collectively, our findings demonstrate that subtype-specific HDTFs in newborn neurons diversify lamina subtypes by repressing default Zfh1-driven identities and promoting alternative fates (Figure 3C). These data establish the newborn-neuron stage as a pivotal window for neuron subtype specification.

### Loss of subtype-specific HDTFs drives subtype-to-subtype fate conversions at molecular, morphological, and functional levels

Loss of Dve converts L2 neurons into L3 based on molecular markers, prompting us to test whether this conversion also encompasses both morphological and functional characteristics. To examine morphology, we genetically labeled L3 neurons with membrane-targeted tdTomato. In adult controls, a single L3 neuron localized to each cartridge edge, extended neurites toward the cartridge core, and projected axons to medulla layer M3 (Figure 4A). In contrast, a single L2 neuron occupied the cartridge center and projects to medulla layer M2^19,20^. In Dve-KD animals, L3 neuron number nearly doubled, with each cartridge containing one endogenous and one ectopic L3 neuron (Figure 4A). Both endogenous and ectopic L3 neurons in Dve-KD occupied cartridge edges, extended neurites into the core, and projected axons to M3, indicating acquisition of canonical L3 morphology and loss of L2 morphology (Figure 4A). Overall axonal targeting remained intact, but terminals in Dve-KD animals appeared broader, with occasional fine neurites extending deeper (Figure 4A), likely due to spatial constraints imposed by the doubled L3 population.

**Figure 4.**
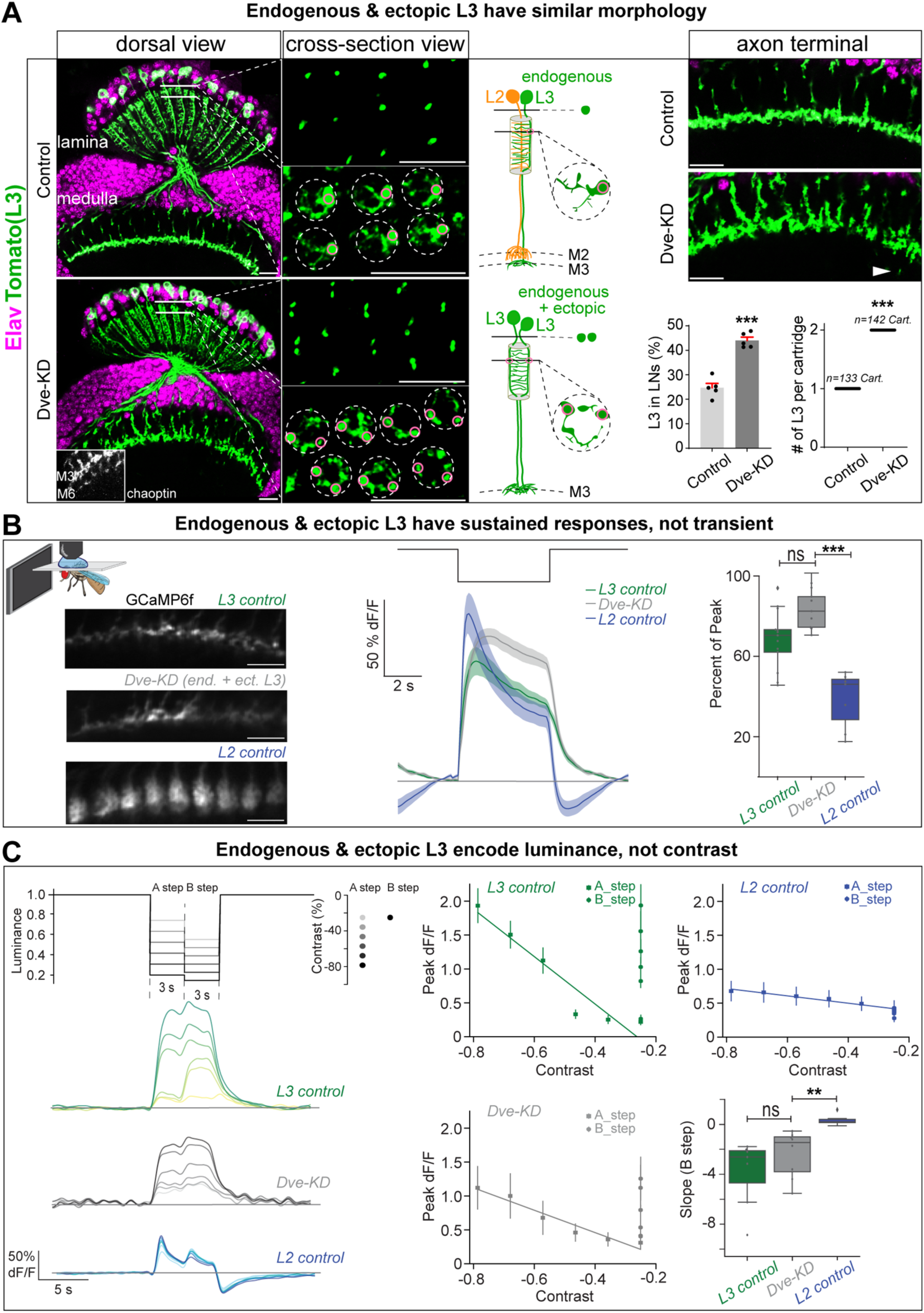
L2-to-L3 converted neurons adopt L3-like morphology and function. (A) L2-to-L3 converted neurons in Dve-KD adopt L3-like dendritic arbors and axonal targeting. Each cartridge contains one endogenous and one ectopic L3 in Dve-KD. Dve-KD was driven by 27G05-Gal4 > UAS-Dve-dsRNA. L3 neurons are labeled with 22E09-LexA > LexAop-myr::tdTomato. Chaoptin labels all photoreceptors and delineates medulla layers M3 and M6. Dashed circles: individual lamina cartridges; pink circles: L3 neurite shafts. The percentage of L3 neurons within total LNs is quantified. Data are mean ± SEM; n = 5 brains; \*\*\**p*<0.001, unpaired t-test. Number of L3 neurons per cartridge is also quantified (control: n = 133 cartridges from 6 brains; Dve-KD: n = 142 cartridges from 6 brains); \*\*\**p*<0.001, permutation test. Scale bar, 10 µm; sample age: 1d adult. (B) L2-to-L3 converted neurons in Dve-KD exhibit L3-like sustained responses rather than L2 transient responses. Dve-KD was driven by 27G05-Gal4 > UAS-Dve-dsRNA. GCaMP6f was expressed in L3 with 22E09-LexA and in L2 with 16H03-LexA. Representative images show GCaMP6f-labeled L3 axonal terminals in control and Dve-KD, and L2 axonal terminals. Scale bar, 10 µm. Average calcium traces were recorded from L3 axonal terminals in response to full-field flashes (FFF) for L3 control (green), Dve-KD (grey), and L2 control (blue). Quantification of the percentage of peak response retained at the OFF-step plateau during FFF. L3 control, n = 13 (297 ROIs); Dve-KD, n = 9 (381 ROIs); L2 control, n = 7 (465 ROIs); ns, not significant, \*\*\**p*<0.001, unpaired t-test. Sample age: 5-7 d adult. (C) L2-to-L3 converted neurons in Dve-KD exhibit L3-like luminance sensitivity rather than L2 contrast sensitivity. Schematic of the A-B steps stimulus paradigm, showing sequential A– and B-steps following a 30 s adapting stimulus. Average calcium traces are shown for L3 control (n = 7, 49 ROIs; green), Dve-KD (n = 8, 37 ROIs; grey), and L2 control (n = 6, 80 ROIs; blue). Peak calcium responses to A– and B-steps are plotted as a function of contrast. Linear regression models were fitted to assess response trends. Slopes from the regression models for peak responses to B-steps are quantified for all conditions. ns, not significant, \*\**p*< 0.01, unpaired t-test. sample age: 5-7 d adult. See also Figure S5 and Movies S1-S3.

We next asked whether these ectopic L3 neurons also acquire L3 functional properties. Although both L2 and L3 neurons respond to light decrements, L2 responses are transient and contrast-sensitive, whereas L3 responses are sustained and luminance-sensitive^21–23^. To assess temporal profiles, we expressed the genetically encoded calcium indicator GCaMP6f in L3 neurons and recorded calcium signals from their axon terminals *in vivo* using two-photon imaging during visual stimulation. In response to a 5-second full-field flash (100% contrast), control L3 neurons showed sustained responses, whereas L2 neurons showed transient responses (Figure 4B; Movie S1 and S3). Upon Dve-KD, all labelled L3 neurons exhibited sustained responses similar to L3 controls (Figure 4B; Movie S1 and S2). Because each medulla column in Dve-KD animals contains both an endogenous and an ectopic L3 neuron, we placed regions of interest (ROIs) on individual axon terminal branches within individual columns to capture single-neuron responses from each. All ROIs responded robustly to the stimulus, with sustained profiles indistinguishable from control L3 neurons, showing that ectopic L3 neurons adopt the L3 sustained response property (Figure S5A).

We then tested luminance sensitivity, a defining property of L3 physiology^21,23^. We used a luminance-contrast discrimination stimulus^23^ (Figure 4C) in which A steps varied in both luminance and contrast, and B steps were fixed at 25% contrast but varied in luminance. L2 neurons responded transiently and equally to all B steps, consistent with contrast sensitivity (Figure 4C). In contrast, all L3 neurons—both endogenous and ectopic—in Dve-KD animals exhibited sustained responses whose amplitudes scaled with luminance, matching the luminance sensitivity of control L3 neurons but distinct from the contrast sensitivity of L2 neurons (Figure 4C). Finally, using a randomized 11-level luminance stimulus, we found that all L3 neurons in Dve-KD exhibited nonlinear encoding of luminance information, consistent with the response property of control L3 neurons (Figure S5B).

Taken together, these data demonstrate that loss of Dve converts L2 to L3 neurons at molecular, morphological, and functional levels, and that the converted neurons are fully integrated into visual circuitry, displaying canonical L3 temporal and luminance-sensitive response properties.

### Pan-LN and subtype-specific HDTFs act combinatorially to specify lamina neuron subtypes

Co-expression of a pan-LN HDTF with subtype-specific HDTFs in newborn neurons suggested that the progenitor-derived pan-LN HDTF could act as a required cofactor to specify distinct subtypes, thereby driving diversification within the progenitor-defined lamina neuron class. We used L5, a genetically tractable subtype, as a representative to test this hypothesis. L5 specification requires the subtype-specific HDTF Bsh; loss of Bsh converts L5 neurons to L1, establishing its necessity for L5 fate^14^. To test whether the pan-LN HDTF Scro cooperates with Bsh, we selectively knocked down Scro in L5 neurons after the onset of Bsh expression, thus keeping Bsh expression intact while depleting Scro during the newborn-neuron stage.

In controls, GFP-labeled L5 neurons co-expressed Scro and Bsh from the newborn-neuron stage, segregated into a discrete soma layer, and subsequently activated the L5-specific HDTF Pdm3 (Figure 5A). In contrast, newborn Scro-knockdown (Scro-KD) neurons transiently expressed Scro prior to RNAi-mediated depletion, retained Bsh, but failed to activate Pdm3 (Figure 5A). These neurons formed disorganized soma clusters intermingled with neighboring L4 cell bodies rather than segregating into the characteristic L5 layer, indicating that Scro is essential for initiating L5-specific marker expression and proper cell body positioning (Figure 5A and 5B).

**Figure 5.**
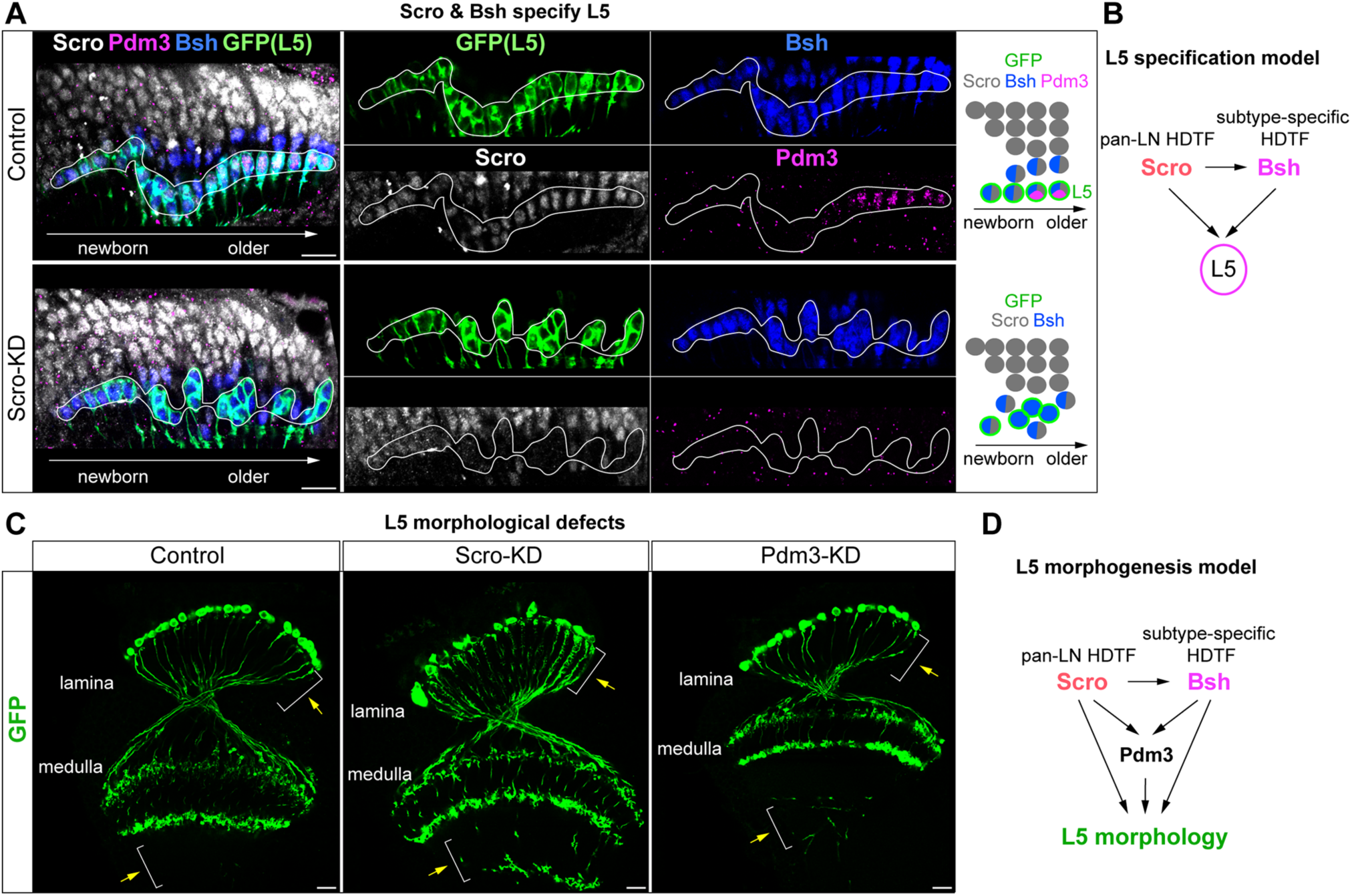
Pan-LN HDTF Scro and subtype-specific HDTF Bsh act combinatorially in newborn neurons to specify L5 neurons. (A) Newborn L5-specific Scro knockdown (Scro-KD: Bsh-L-Gal4 > UAS-Scro-RNAi) yields transient Scro expression prior to RNAi-mediated depletion. Bsh remains intact, but Pdm3 induction is blocked. Bsh-LexA > LexAop-myr::GFP labels L5 neurons independently of fate. Pdm3: L5-specific HDTF. White outlines: GFP^+^ neurons. Sample age: 19h APF. (B) Schematic model: Scro and Bsh cooperate to specify L5 neurons. Our previous work shows that Bsh is required in newborn neurons to specify L5 identity^14^. (C) Scro-KD neurons exhibit ectopic neurites in the lamina and axonal misprojection in the medulla. Pdm3-KD (Bsh-L-Gal4 > UAS-Pdm3-RNAi-#1) partially phenocopies the medulla projection defect. Yellow arrows highlight ectopic neurites in the lamina (Scro-KD) and misprojected axons in the medulla (Scro-KD and Pdm3-KD). Sample age: 1d adult. (D) Schematic summary: Scro and Bsh act combinatorially to specify L5 morphology, in part via Pdm3. Our previous work shows that Bsh is required for L5-specific morphology^14^. Scale bar, 10 µm; n ≥ 5 brains. See also Figure S6.

In adults, Scro-KD neurons did not acquire characteristic L5 morphological features. In controls, L5 neurons lacked neurites spanning the lamina and projected axons into two superficial medulla layers (M1 and M5), whereas Scro-KD neurons extended ectopic neurites throughout the lamina and projected aberrant axonal extensions into deeper medulla layers (Figure 5C and S6). Notably, knocking down Pdm3 alone generated the axonal misprojection phenotype but did not disrupt soma organization or induce ectopic neurite extension within the lamina (Figure 5C and S6). This observation suggests that Scro regulates additional downstream targets beyond Pdm3 that are required for complete L5 morphogenesis (Figure 5D).

Together, these findings demonstrate that Scro and Bsh act cooperatively at the newborn-neuron stage to specify L5 neuron identity. More broadly, they support a model in which progenitor-derived pan-LN HDTF induces subtype-specific HDTFs and functions as their obligate cofactor in newborn neurons to specify distinct subtypes, thereby driving diversification within a progenitor-defined lamina neuron class (Figure 2B and 5B).

### Mouse retina: conservation of pan-class and subtype-specific HDTF logic

To determine whether the progenitor-derived regulatory logic for within-class subtype diversification is conserved from *Drosophila* to mammals, we examined HDTF expression patterns in mouse retinal neurons. Between embryonic day 11 and postnatal day 7, retinal progenitor cells generate six major retinal neuron classes—retinal ganglion cells (RGCs), cones, amacrine cells (ACs), horizontal cells (HCs), rods, and bipolar cells (BCs) (Figure 6A). During this neurogenic period, retinal progenitor cells receive Sonic Hedgehog (Shh) signaling^24,25^, suggesting that, analogous to *Drosophila*, Shh may initiate HDTF cascades essential for neuronal diversification.

**Figure 6.**
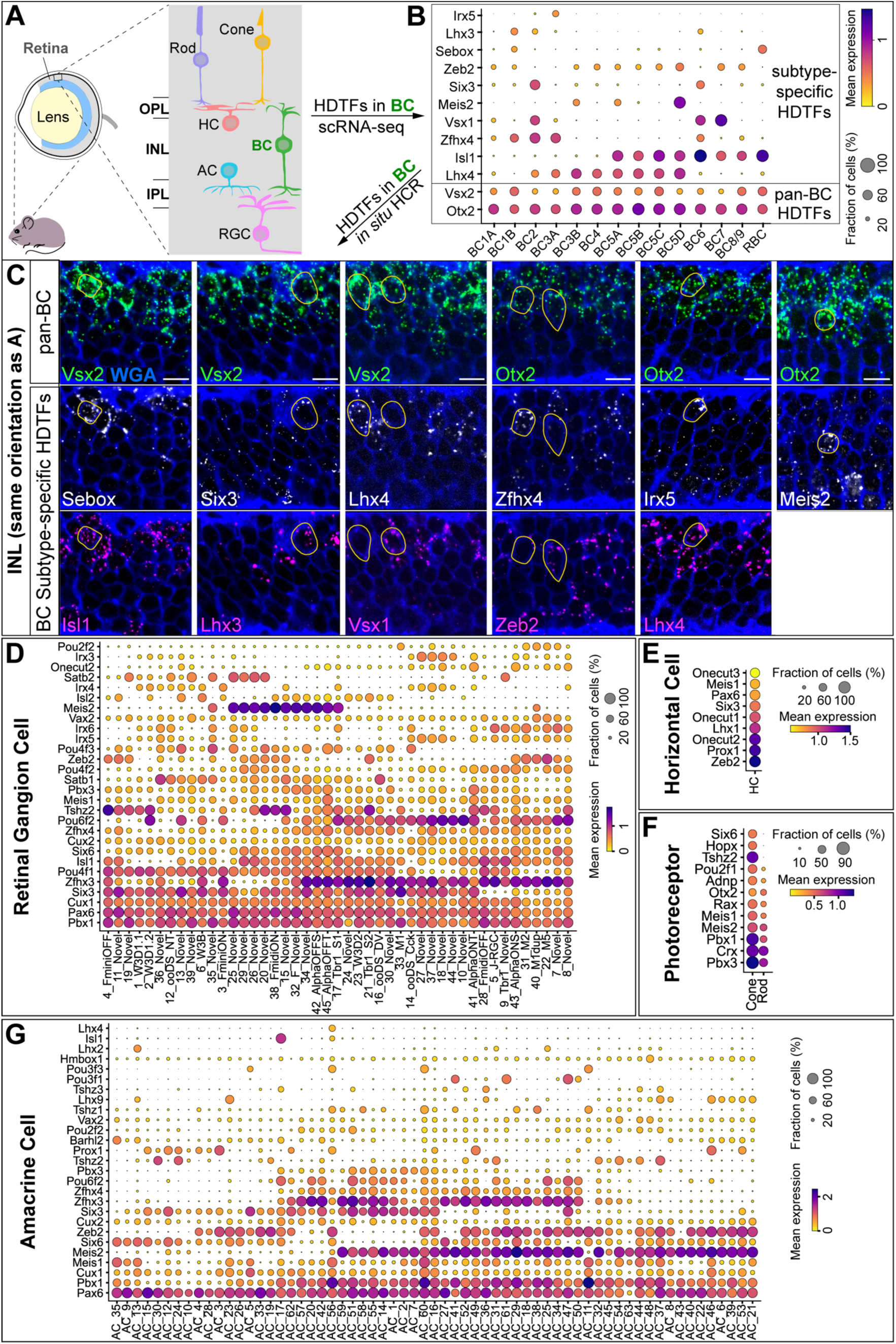
Mouse retinal neurons express pan-class and subtype-specific HDTFs. (A) Schematic of the mouse retina. BC: bipolar cell; HC: horizontal cell; RGC: retinal ganglion cell; AC: amacrine cell. (B) HDTF expression across 15 BC subtypes from published scRNA-seq data^26^ (P17). Dot size indicates the percentage of cells in each subtype expressing a given gene; color indicates mean normalized expression among expressing cells. (C) Validation of pan-BC and subtype-specific HDTF expression in P18 retinas by *in situ* hybridization chain reaction (HCR). Yellow circles: subtype-specific HDTF^+^ BC somata. (D-G) HDTF expression in cones, rods, horizontal cells, retinal ganglion cells, and amacrine cells from a single-cell atlas of the adult mouse retina^2^. Scale bar, 10 µm; n = 3 mice. See also Supplementary Data 1.

We first asked whether BCs co-express pan-class and subtype-specific HDTFs. Analysis of published scRNA-seq data^26^ from mature BCs revealed that all 15 BC subtypes uniformly express the pan-BC HDTFs Otx2 and Vsx2^27^ (Figure 6B). Consistent with our *Drosophila* findings, these pan-BC HDTFs are initiated in progenitors^27^. By contrast, BC subtypes differentially expressed distinct sets of subtype-specific HDTFs, including established subtype markers Lhx3, Lhx4, Meis2, Irx5, Vsx1, Isl1, and Zeb2^27–32^ (Figure 6B). Using *in situ* hybridization chain reaction (HCR) on postnatal day 18 retinas, we validated the differential expression of these subtype-specific HDTFs within subsets of Vsx2+ or Otx2+ BCs (Figure 6C; Supplementary Data 1). We also identified previously unreported HDTFs—Sebox, Six3, and Zfhx4—in subsets of Vsx+ or Otx2+ BC neurons in the inner nuclear layer (INL) (Figure 6C). Although Vsx2 is also expressed in Müller glia, these populations occupy distinct INL regions, with BC cell bodies located near the outer plexiform layer (OPL) and Müller glia situated centrally^28^. Notably, Sebox and Six3 transcripts localized to Vsx2+ cells adjacent to the OPL, confirming their BC-specific expression (Figure 6C). Together, these data show that BCs co-express pan-BC HDTFs, initiated in progenitors, alongside subtype-specific HDTFs.

To assess whether this HDTF expression logic extends to other retinal neuron types, we analyzed a recent single-cell atlas of the adult mouse retina^2^. All major retinal neuron types—cones, rods, HCs, RGCs, and ACs—expressed distinct combinations of HDTFs (Figure 6D–6G). Notably, RGCs and ACs, both highly diversified (45 and 63 subtypes, respectively)^2^, co-expressed candidate pan-class HDTFs together with subtype-specific HDTFs (Figure 6D and 6G).

Together, these findings demonstrate that mouse retinal neurons, like *Drosophila* lamina neurons, co-express pan-class and subtype-specific HDTFs, indicating an evolutionarily conserved pan-class/subtype-specific HDTF logic (Figure 7).

**Figure 7.**
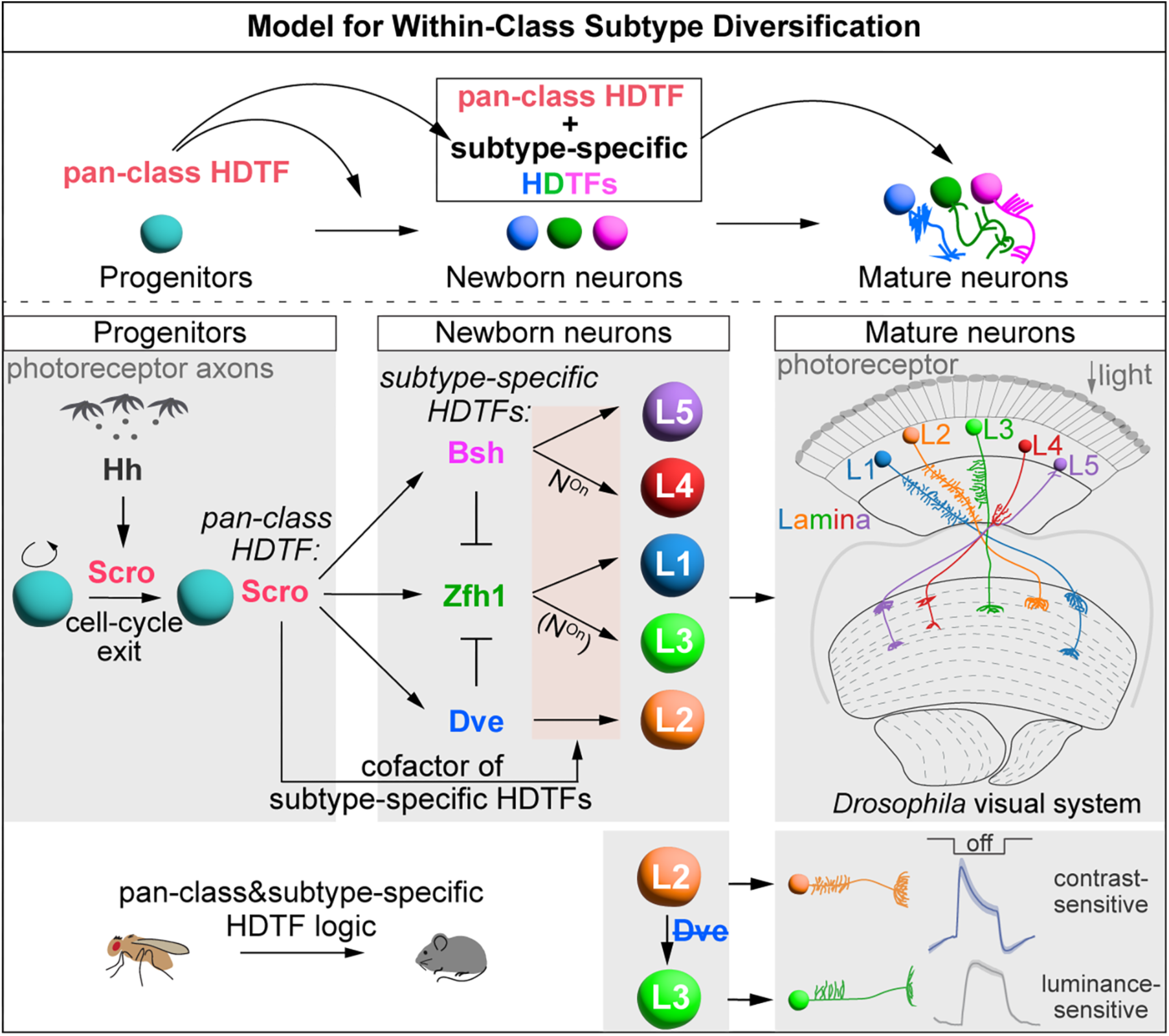
Model for within-class subtype diversification. We propose a generalizable progenitor-derived regulatory logic: a pan-class HDTF induced in progenitors persists in newborn neurons, induces subtype-specific HDTFs, and acts as their obligate cofactor to specify distinct subtypes. In the *Drosophila* lamina, photoreceptor-derived Hedgehog (Hh) induces the pan-class HDTF Scro in progenitors, promoting cell-cycle exit; Scro is inherited by all newborn lamina neurons, where it activates subtype-specific HDTFs (Dve, Zfh1, Bsh) and acts as their cofactor to specify distinct lamina neuron subtypes. In the mouse retina, neurons exhibit pan-class and subtype-specific HDTF expression, indicating evolutionary conservation of this logic. N^on^: Notch^on^; (N^on^): predicted Notch^on^.

## DISCUSSION

### Hedgehog Signaling and Neural Progenitor Identity

Neural progenitors respond to extrinsic or intrinsic cues to produce diverse neuron types. For example, in the *Drosophila* embryonic ventral nerve cord, secreted Wingless and Hh proteins specify neuroblast identities^33–36^. Similarly, in the mouse ventral neural tube, graded Shh signaling defines distinct neural progenitor domains by directing combinatorial HDTF expression^37,38^. Shh signaling also drives the production of dopaminergic^39^ and serotonergic neurons^40^ in the mouse midbrain and hindbrain, respectively, likely by defining progenitor identities.

Previous work showed that Hh secreted from photoreceptors is required for LPCs to undergo terminal division^12^. Here, we extend these findings by demonstrating that Hh signaling activates the pan-LN HDTF Scro in LPCs, promoting cell-cycle exit and initiating expression of subtype-specific HDTFs—Zfh1, Dve, and Bsh—in newborn neurons to drive lamina neuron subtype diversification. Whether this Hh-dependent HDTF cascade is conserved in other neural systems remains an important question. Notably, in the mouse retina, progenitors receive Shh signals from early-born retinal ganglion cells (RGCs)^41^, and retinal neurons exhibit a similar pan-class and subtype-specific HDTF expression pattern, supporting evolutionary conservation. Given that retinal progenitors and other neural progenitors sequentially express temporal TFs, future studies should investigate how Hh signals integrate with temporal TF programs to generate neuronal diversity.

### Newborn Neurons: A Critical Window for Neuronal Diversification

Temporal transcriptomic profiling across neuronal development—from progenitors to mature neurons—in systems such as the *Drosophila* visual system, mouse retina, neocortex, and somatosensory neurons has revealed the progressive emergence of transcriptionally distinct neuron types and identified candidate regulators of this diversification^5,42–44^. However, most analyses have focused on developing^5,42–47^ or mature neurons^48,49^, rather than newborn neurons, emphasizing how their differentially expressed genes contribute to morphological and functional specialization. Newborn neurons have received limited attention, partly because their transcriptomes and morphology appear uniform—they have not yet developed dendrites, axons, or synapses.

Despite their apparent uniformity, subtle molecular differences among newborn neurons can critically influence their subsequent differentiation. For instance, transient Notch signaling in newborn neurons governs binary fate decisions^50–52^. In the *Drosophila* lamina, we find that, in newborn neurons, progenitor-derived pan-class HDTF both induces subtype-specific HDTFs and acts as their required partner to specify distinct lamina neuron subtypes. Loss of subtype-specific HDTFs during the newborn-neuron stage—but not at later stages—causes subtype-specific fate conversions at molecular, morphological, and functional levels, underscoring a temporally restricted requirement for subtype-specific HDTFs in subtype specification. Our previous work^53^ further shows that Notch signaling integrates with HDTF activity to diversify neuronal fates (Figure 7). Defining the molecular logic that integrates Notch signaling with HDTF activity in newborn neurons will be critical. Together, these findings highlight the newborn-neuron stage as a pivotal window in which progenitor-derived programs are interpreted, thereby determining future gene-expression profiles and functional properties of mature neurons.

### Within-Class Subtype Diversification by Pan-class and Subtype-Specific HDTFs

While spatial factors and temporal TFs endow progenitors with the capacity to produce diverse neuron types^54,55^, a key gap remains: how can progenitors generate multiple, functionally related subtypes within a single neuronal class? Here, in *Drosophila*, we identify an HDTF cascade acting across the progenitor-to-newborn neuron transition that drives subtype diversification within a progenitor-defined class. We also uncover an analogous pan-class/subtype-specific HDTF expression logic in the mouse retina. For instance, each of 45 RGC subtypes expresses candidate pan-RGC HDTFs together with subtype-specific HDTFs, consistent with a conserved within-class diversification strategy.

Several fundamental questions arise. Do pan-class HDTFs originate in retinal progenitor cells, and which upstream signals initiate their expression? Do pan-class HDTFs activate subtype-specific HDTFs in newborn neurons, as seen for Scro in the lamina? Are both pan-class and subtype-specific HDTFs required to specify individual RGC subtypes? And finally, do subtype-specific HDTFs diversify RGCs from a common transcriptional ground state?

Our *Drosophila* data suggest a model in which the pan-LN HDTF Scro initiates neuronal diversification by selectively activating Zfh1, Dve, and Bsh in distinct newborn neuron populations. The basis of this selectivity is an important open question. The ectopic expression of Zfh1 upon Dve or Bsh knockdown suggests that these subtype-specific HDTFs repress Zfh1. This repression could occur either through direct transcriptional repression by Dve and Bsh or via long-range chromatin interactions, akin to regulatory mechanisms governing Hox gene clusters and olfactory receptor loci^56–58^.

Given the brain-wide expression of HDTFs across species^9–11^, our findings define a generalizable progenitor-to-newborn neuron HDTF cascade by which progenitor identity directs subtype diversification within a neuronal class, with direct implications for subtype-precise reprogramming and cell-replacement therapies (e.g., restoring retinal ganglion cell subtypes lost in glaucoma).

## RESOURCE AVAILABILITY

### Lead contact

Further information and requests for resources and reagents should be directed to, and will be fulfilled by, the lead contact, Chris Doe.

### Materials availability

Further information and requests for resources and reagents should be directed to the lead contact.

### Data and code availability

The RNA-seq data reported in this paper have been deposited at GEO under accession number GSE305551 and are publicly available. All original code has been deposited at [www.github.com/pnewstein/multi-species-homeodomain] and is publicly available.

## ACKNOWLEDGMENTS

We thank Lawrence Zipursky for sharing published and unpublished reagents. We thank Claude Desplan, Makoto Soto, Richard Mann, James Skeath, Jing Peng, and Cheng-Ting Chien for reagents. Antibodies obtained from the Developmental Studies Hybridoma Bank, created by the NICHD of the NIH and maintained at the University of Iowa, Department of Biology, Iowa City, IA, were used in this study. Stocks obtained from the Bloomington Drosophila Stock Center (NIH P40OD018537) and Vienna Drosophila Resource Center were used in this study. We thank Ben Brissette, Kristen Lee, and Minoree Kohwi for comments on the manuscript.

This article is subject to HHMI’s Open Access to Publications policy. HHMI lab heads have previously granted a nonexclusive CC BY 4.0 license to the public and a sublicensable license to HHMI in their research articles. Pursuant to those licenses, the author-accepted manuscript of this article can be made freely available under a CC BY4.0 license immediately upon publication.

## AUTHOR CONTRIBUTIONS

CX conceived the project, performed and analyzed all experiments except neuron function assays, and wrote the manuscript with feedback from all authors. RGS performed and analyzed experiments. PN analyzed scRNA-seq data and designed HCR probes. RND helped perform mouse experiments. NW and MS performed and analyzed neuron function assays and contributed to the associated figures and text. CLC provided feedback on mouse experiments. CQD provided feedback during the project, contributed to writing the paper and assembling the figures. Funding was provided by HHMI (CQD, CLC) and from the European Research Council through the ERC grant ‘Adaptive Vision’, agreement No. 101045003 (MS). All authors commented on and approved the final manuscript.

## DECLARATION OF INTEREST

The authors declare no competing financial or non-financial interests.

**Supplementary Data 1. Custom mouse HCR probe sets and their sequences, related to Figure 6**.

**Movie S1. GCaMP6f calcium responses in L3 neurons of control animals during full-field visual stimulation, related to Figure 4**. This movie corresponds to Figure 4B. Frames were acquired at ∼10 Hz. Alternating full-field OFF (0 × Imax) and ON (Imax) flashes, each lasting 5 seconds, were delivered with stimulation onset at 0.0 seconds, 10.0 seconds, and so on. Playback speed is real time. Sample age: 5-7 d adult.

**Movie S2. GCaMP6f calcium responses in L3 neurons of Dve-KD animals during full-field visual stimulation, related to Figure 4**. This movie corresponds to Figure 4B. Frames were acquired at ∼10 Hz. Alternating full-field OFF (0 × Imax) and ON (Imax) flashes, each lasting 5 seconds, were delivered with stimulation onset at 0.0 seconds, 10.0 seconds, and so on. Playback speed is real time. Sample age: 5-7 d adult.

**Movie S3. GCaMP6f calcium responses in L2 neurons during full-field visual stimulation, related to Figure 4**. This movie corresponds to Figure 4B. Frames were acquired at ∼10 Hz. Alternating full-field OFF (0 × Imax) and ON (Imax) flashes, each lasting 5 seconds, were delivered with stimulation onset at 0.0 seconds, 10.0 seconds, and so on. Playback speed is real time. Sample age: 5-7 d adult.

## METHOD

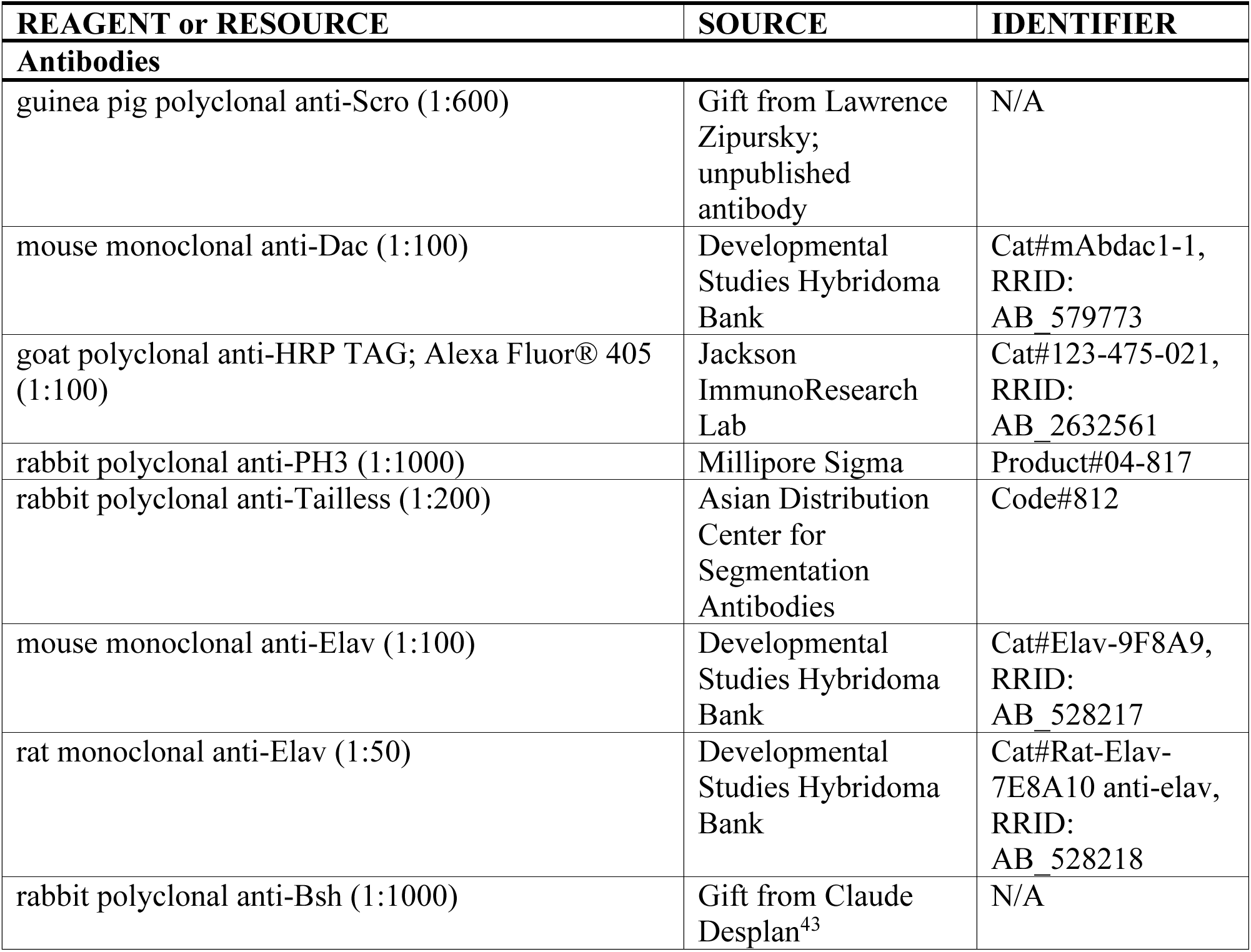
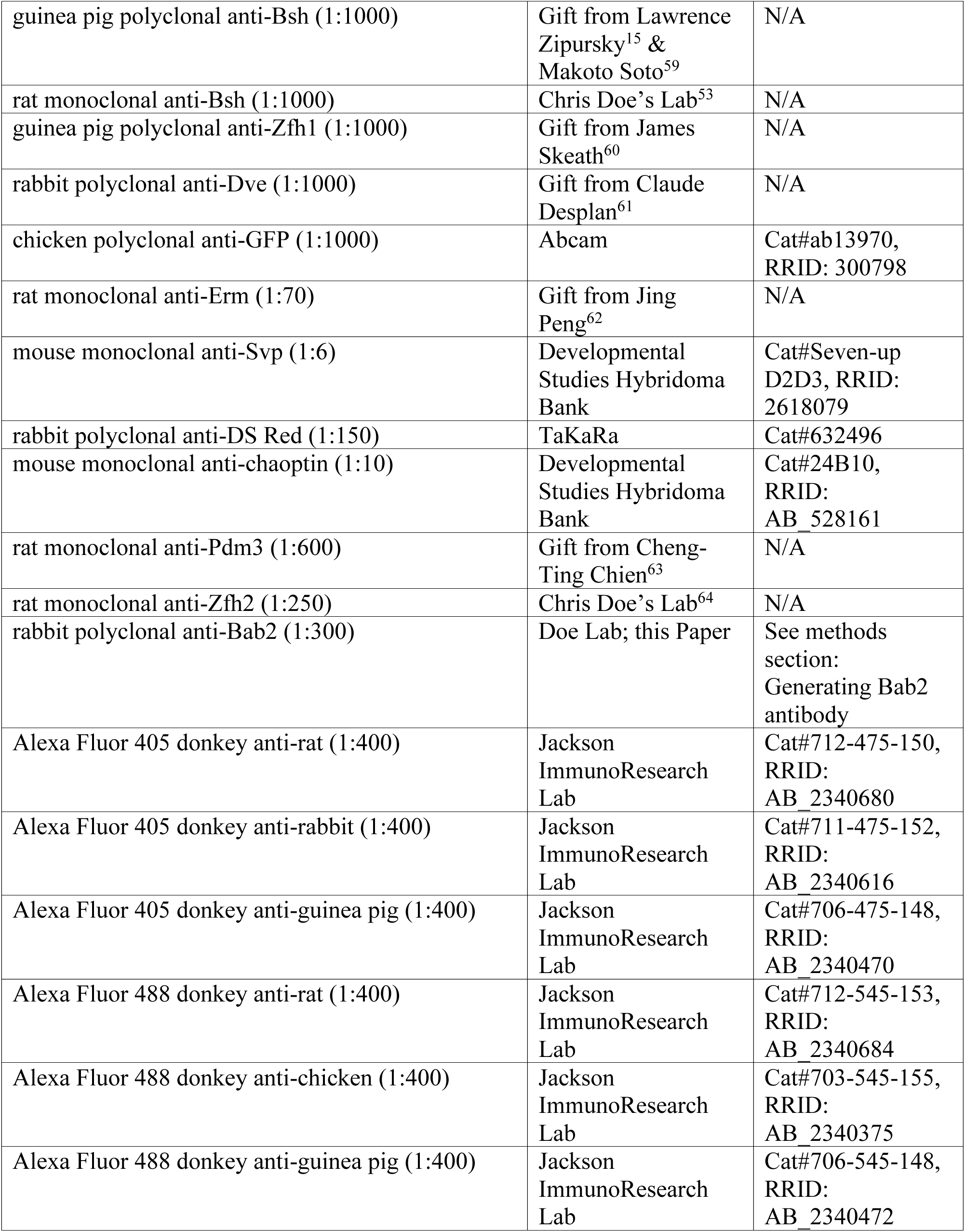
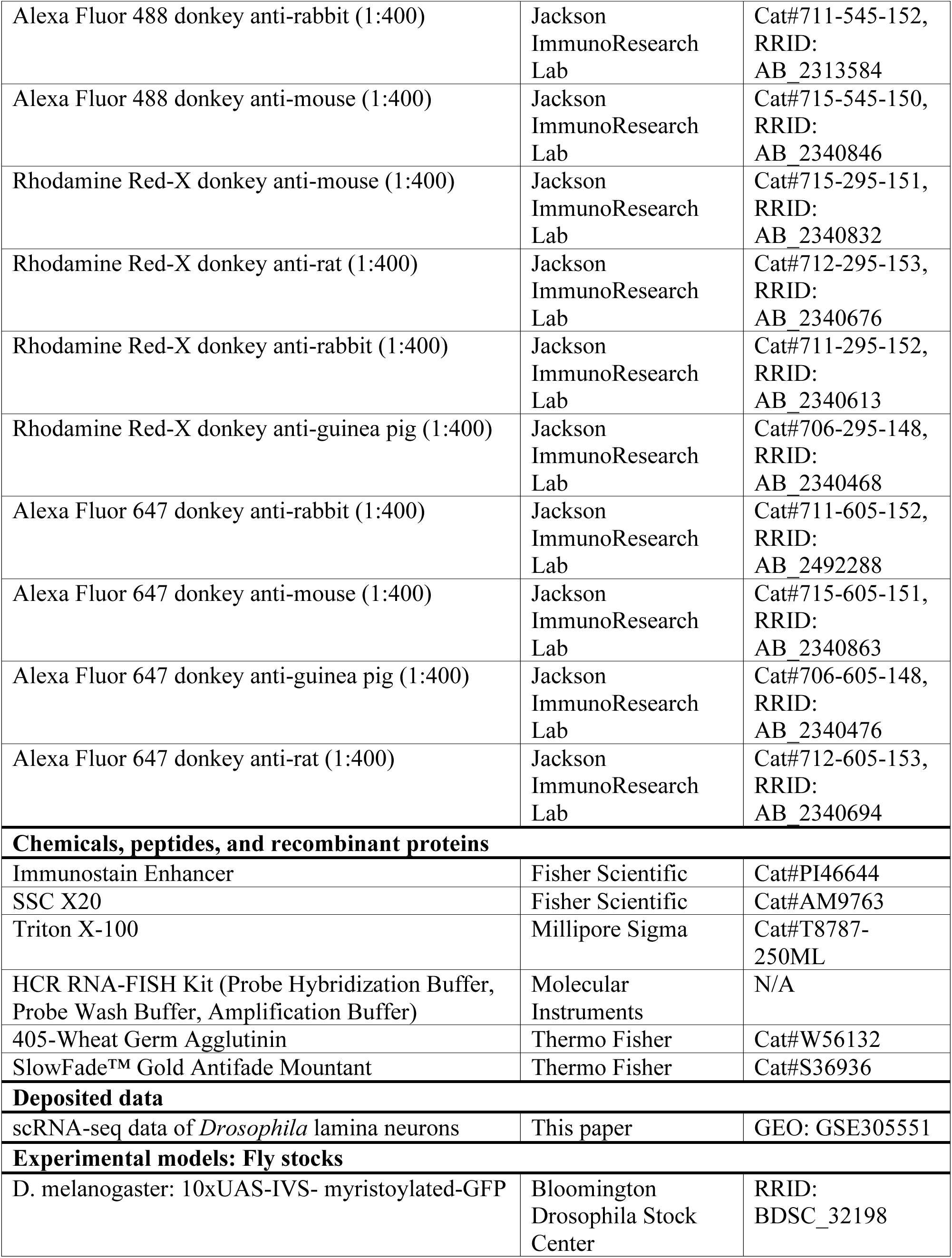
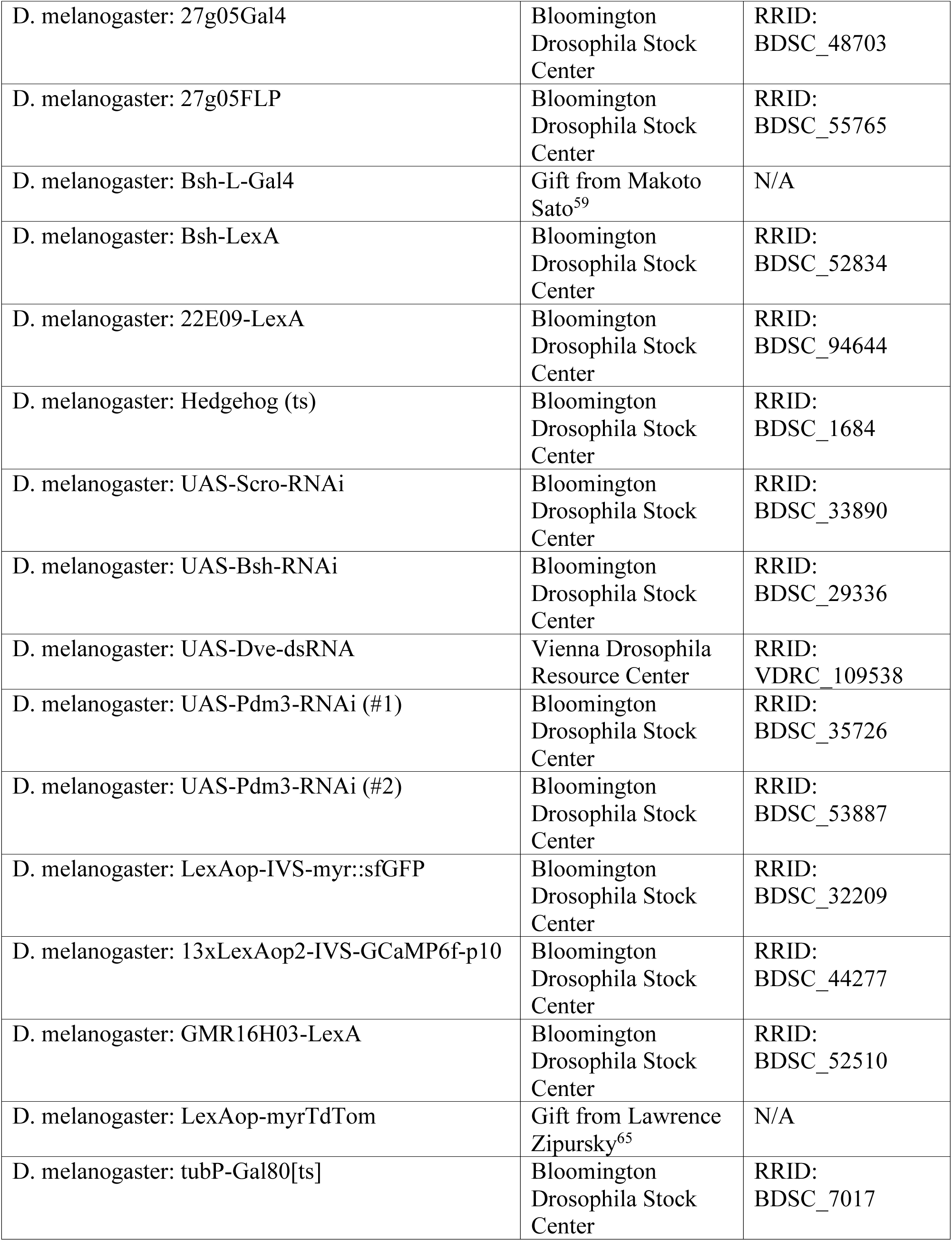
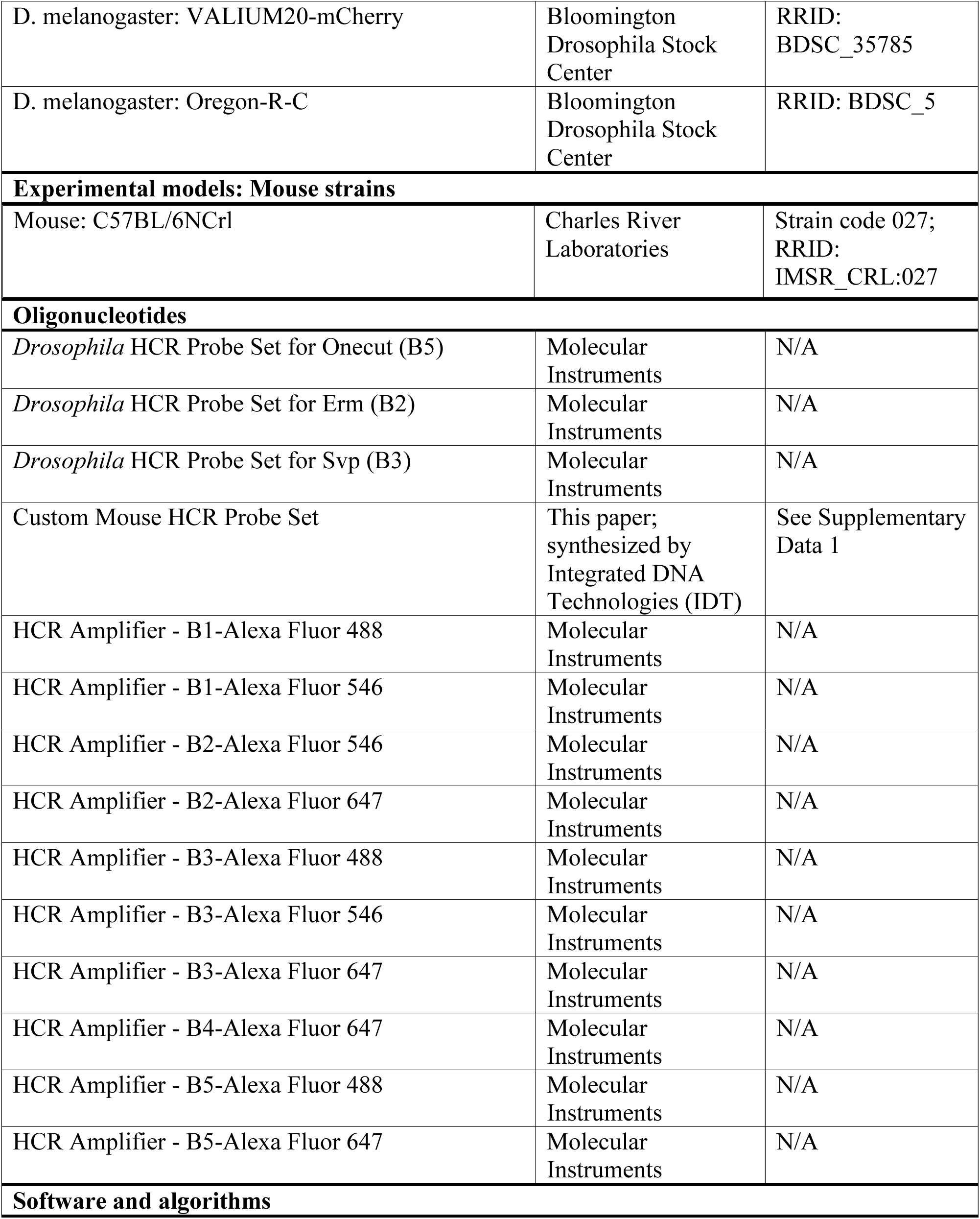
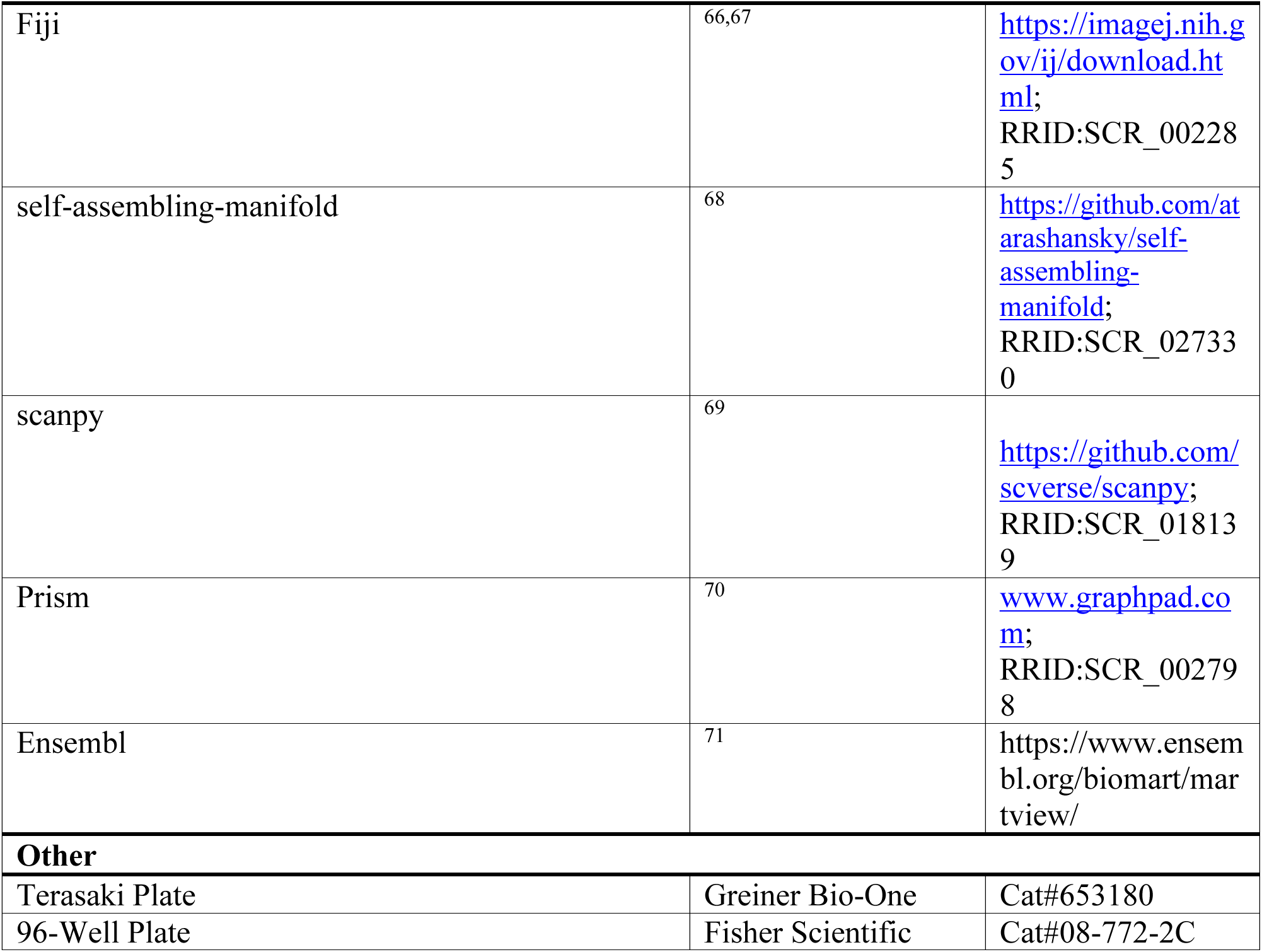
KEY RESOURCES TABLE.

## EXPERIMENTAL MODEL AND SUBJECT DETAILS

All flies were reared at 25 °C on standard cornmeal fly food, unless otherwise stated. For all RNAi and dsRNA knockdown experiments, crosses were kept at 25 °C, and their progeny were kept at 28°C with a 16:8 hours light-dark cycle from the second larval stage until dissection. For all Gal80^ts^ experiments, crosses were kept at 18°C, and progeny were kept at 29 °C at the desired time.

All mouse experiments were conducted in compliance with the protocol IS00001679, approved by the Institutional Care and Use Committees (IACUC) at Harvard University. Experiments were performed on tissue collected from wild-type male and female C57BL/6NCrl mice (Charles River). Tissues were collected on postnatal day (P) 18 for all mouse experiments.

## METHOD DETAILS

### Animal collections

#### Hedgehog (Hh) loss-of-function

For the temperature-sensitive Hh loss-of-function experiment (*hh*^ts^), animals were reared at 18 °C and shifted to 28 °C at the onset of wandering third-instar larvae (L3). Brains were dissected at 19 h after pupal formation (APF). *hh*^ts^/TM6B served as controls; *hh*^ts^/*hh*^ts^ were the loss-of-function genotype.

#### Conditional Scro knockdown in LPCs and progeny

For Scro-knockdown (Scro-KD) using 27G05-Gal4, tub-Gal80^ts^, and UAS-Scro-RNAi, animals were reared at 18 °C and shifted to 29 °C at early L3 to permit Gal4-driven RNAi expression and initiate Scro depletion just before the lamina neurogenesis.

#### Bsh-KD, Dve-KD, and Bsh/Dve-DKD

For Bsh-KD, Dve-KD, and Bsh/Dve double knockdown (DKD) using 27G05-Gal4 with UAS-Bsh-RNAi and UAS-Dve-dsRNA, animals were reared at 25 °C and shifted to 28 °C at the second-instar larval stage.

#### Scro-KD in newborn L5 neurons

For Scro-KD with Bsh-L-Gal4 and UAS-Scro-RNAi, animals were reared at 25 °C and shifted to 29°C at early L3.

#### scRNA-seq samples

Oregon-R flies were reared at 25 °C, and 3-day-old adult females were dissected for scRNA-seq.

### Generating Bab2 antibody

Bab2 protein and antibody production were performed by GenScript (Piscataway, NJ) using rabbits immunized with the following portion of the Bab2 open reading frame:

RDPEHT ELRMCLEAKK SRSLPVSPQP QPNLKLAGSA LFEFGQRSSP VETKIKTNPE TKPPRRKIVP PSGEGQQFCL RWNNYQSNLT NVFDELLQSE SFVDVTLSCE GHSIKAHKMV LSACSPYFQA LFYDNPCQHP IIIMRDVSWS DLKALVEFMY KGEINVCQDQ INPLLKVAET LKIRGLAEVS AGRGEGGASA LPMSAFDDED EEEELASATA ILQQDGDADP DEEMKAKRPR LLPEGVLDLN QRQRKRSRDG SYATPSPSLQ GGESEISERG SSGTPGQSQS QPLAMTTSTI VRNPFASPNP QTLEGRNSAM NAVANQRKSP APTATGHSNG NSGAAMHSPP GGVAVQSALP PHMAAIVPPP PSAMHHHAQQ LAAQHQLAHS HAMASALAAA AAGAGAAGAG GAGSGSGSGA SAPTGGTGVA GSGAGAAVG

### *Drosophila* Immunohistochemistry

Fly brains were dissected in Schneider’s medium and fixed in 4% paraformaldehyde (PFA) diluted in phosphate-buffered saline (PBS) for 25 min. Following fixation, brains were briefly rinsed in PBS containing 0.5% Triton X-100 (PBST) and then incubated in PBST for ≥4 hr at room temperature (RT) in a 96-well plate. Samples were subsequently blocked overnight (O/N) at 4 °C in blocking buffer (10% normal donkey serum, PBST). Brains were then incubated in primary antibody (diluted in immunostain enhancer solution) at 4 °C for ≥2 nights. After four 25-min washes with PBST, samples were incubated in secondary antibody (diluted in blocking buffer) at 4 °C for ≥1 day. After secondary antibody incubation, brains were washed with PBST four times for 25 min each and mounted in SlowFade Gold antifade reagent. Images were acquired using a Zeiss 800 confocal and processed with Image J^66^.

### *Drosophila* HCR-FISH

The HCR-FISH protocol was adapted from Dombrovski et al.^72^ with minor modifications. Following dissection and fixation on ice, samples were rinsed three times with PBST and incubated in PBST for 30 min on ice in a Terasaki plate. Samples were transferred to a 96-well plate and equilibrated in pre-warmed probe hybridization buffer (PHB) for 30 min at 37 °C and then incubated O/N at 37 °C in probes diluted in PHB (1 μL of 1 μM stock probe solution per 200 μL buffer).

The next day, the samples were washed four times with pre-warmed probe wash buffer (PWB) for 15 min at 37 °C, followed by two 5 min washes with SSCT buffer (20x SSC with 0.1% Triton X-100) rotating at RT. Samples were equilibrated in HCR amplification buffer (AB) for 30 min, rotating at RT. During this time, HCR Amplifiers were heat-activated in a thermal cycler (90 s at 95 °C), and then kept in the dark for 30 min at RT. The HCR Amplifiers were diluted in AB (2 μl of each hairpin per 100 μl of buffer), and incubated the samples at RT O/N.

Following amplification, samples were washed four times with SCCT at RT: twice for 5 min and twice for 30 min. Samples were mounted in SlowFade Gold antifade reagent and stored at 4 °C until confocal imaging using a Zeiss 800 microscope. HCR-FISH was performed using HCR v3.0 probe sets, which were designed and synthesized by Molecular Instruments; probe sequences are proprietary and were not disclosed by the manufacturer.

### *Drosophila* scRNA-seq sample preparation

Single-cell RNA sequencing (scRNA-seq) was performed using a protocol adapted from Peng et al.^73^ with minor modifications. Papain (100 U/ml in Complete Schneider’s Medium, CSM) was freshly prepared and activated at 37 °C for 30 min; Liberase TM aliquots were thawed at RT. Adult brains were dissected in cold CSM, and optic lobes were pooled on ice in Rinaldini’s Solution (RS). For each dissociation, 300 μl activated papain was combined with 4.1 μl Liberase (0.18 WU/ml) to prepare the dissociation solution. After a brief RS wash and removal, the dissociation solution was added to the lobes and incubated at 25 °C with shaking (1000 rpm) for 15 min, with gently mixing at 5 min and 10 min. Lobes were washed twice with RS, followed by two washes with CSM, and then gently triturated in fresh CSM to obtain a homogenous single-cell suspension. Suspensions were passed through a 20 μm strainer into low-adhesion tubes on ice; the filter was rinsed with 500 μl CSM. Cells were pelleted (300 × g, 5 min, RT), resuspended in PBSB (PBS + 0.04% BSA), and submitted immediately for 10x Genomics scRNA-seq. Solutions: CSM (50 mL): 5 mL heat-inactivated bovine serum; 0.1 mL insulin; 5 mL 200 mM L-glutamine; 12.5 µL 20 mg/mL L-glutathione; 39 mL Schneider’s medium; 1 mL penicillin/streptomycin. RS (10× stock, 100 mL): 8 g NaCl; 200 mg KCl; 50 mg NaH₂PO₄; 1 g NaHCO₃; 1 g glucose. (Dilute to 1× before use.)

### *Drosophila* scRNA-seq analysis

Gene-by-cell count matrices were generated with 10x Genomics Cell Ranger (v7.1.0). Quality control retained cells with 500–2,500 detected genes and ≤10% mitochondrial UMI fraction. Downstream preprocessing used the Self-Assembling Manifold (SAM) package^68^ with default parameters for normalization, highly variable gene selection, and dimensionality reduction. Cells were clustered with the Leiden algorithm^74^ (resolution = 1.0), and clusters corresponding to L1–L5 were annotated by canonical marker gene expression. Data were visualized using UMAP as implemented in Scanpy^69^.

### HDTF expression profiling (*Drosophila* and Mouse)

*Drosophila* lamina single-cell data were generated in this study; processing details are provided above. Mouse bipolar-cell scRNA-seq data were obtained from Shekhar et al. ^75^ and converted to AnnData using Scanpy^69^; scRNA-seq data for other retinal classes were obtained from Li et al.^2^. HDTFs were defined by SMART^76^ or PANTHER^77^ domain annotations accessed via Ensembl BioMart^78^. Dot-plot visualizations were produced with Scanpy^69^. Genes were retained if their mean normalized expression in any cell type within a class exceeded a prespecified threshold, and genes were ordered by the number of cell types within that class surpassing the threshold.

### *Drosophila* Lamina neuron physiology

#### Two-photon imaging

Female flies aged 5-7 days post-eclosion were reared on standard molasses-based food under a 12:12 h light-dark cycle at 25 °C and 60% humidity. Two-photon imaging was performed at room temperature (20 °C). Prior to imaging, flies were cold-anesthetized and mounted onto stainless-steel foil with an opening for the thorax and head. Flies were head-fixed using UV-sensitive glue (Bondic), and their legs were immobilized with bee wax to ensure stability. The head was tilted downward, orienting the fly towards the stimulation screen and exposing the cuticle at the back of the head for dissection. A small window was cut in the cuticle at the back of the head using breakable razor blades. During imaging, the brain was perfused with a carboxygenated saline solution containing: 103 mM NaCl, 3 mM KCl, 5 mM TES, 1 mM NaH₂PO₄, 4 mM MgCl₂, 1.5 mM CaCl₂, 10 mM trehalose, 10 mM glucose, 7 mM sucrose, and 26 mM NaHCO₃. For dissections, the same solution was used without calcium and sugars. The pH of the saline equilibrated near 7.3 when bubbled with 95% O₂ / 5% CO₂. Imaging was performed on a Bruker Ultima microscope equipped with a 20×/NA1.0 objective (Leica, Wetzlar, Germany). Excitation was provided by a fixed-wavelength 930 nm laser (YLMO-930, Menlo Systems, Martinsried, Germany), with 5–15 mW power at the sample. Emitted light passed through a SP680 short-pass filter, a 560 lpxr dichroic mirror, and a 525/70 emission filter. Data were acquired using PrairieView software at ∼10 Hz frame rate with an optical zoom of 6×–8×.

#### Visual stimulation for two-photon imaging

All visual stimuli were generated using custom-written software in PsychoPy^79^ and projected onto a 9 × 9 cm rear-projection screen placed at a 45° angle in front of the fly, covering 80° of the visual field in both azimuth and elevation. Stimuli were delivered using a LightCrafter projector (Texas Instruments, Dallas, TX, USA) at a frame rate of 100 Hz. The projected light was filtered through a 482/18 band-pass filter and a ND1.0 neutral density filter (Thorlabs). The maximum luminance (I_max_) at the screen was 4.8 µW cm^-2^.

#### Full-field flashes

The stimulus consisted of alternating full-field ON (I_max_) and OFF (0 × I_max_) flashes, each lasting 5 s. These epochs were repeated at least seven times.

#### 11_steps_luminance

This stimulus consisted of 7 s full-field flashes at 11 luminance levels: 0, 0.1, 0.2, 0.3, 0.4, 0.5, 0.6, 0.7, 0.8, 0.9, and 1 × I_max_. Epochs were randomized and repeated at least three times.

#### A_B_steps

To distinguish contrast-sensitive from luminance-sensitive neurons, we used a stimulus protocol with two consecutive OFF steps (A and B), each lasting 3 s, following a 30 s adapting grey period. The A step consisted of one of six linearly decreasing luminance values, producing six different Weber contrast values relative to the adapting luminance. The subsequent B step also took on one of six decreasing luminance values, selected based on the preceding A step, resulting in six uniform 25% Weber contrast steps. A_B epochs were randomized and repeated at least three times.

#### Data analysis

Data processing was performed using Python 2.7^80^. Pre-processing included motion correction using a Hidden Markov Model from the SIMA Python package^81^, followed by manual selection of regions of interest (ROIs). For each imaging frame, the average intensity within each ROI was calculated and background-subtracted to generate time traces. All neural responses and visual stimuli were interpolated to a temporal resolution of 10 Hz. Neural activity was quantified as relative changes in fluorescence over time (ΔF/F₀), where F₀ was defined as the mean fluorescence across the entire trace. Analyses included trial averaging and stimulus-specific evaluations, performed per ROI and then averaged across flies. ROIs showing responses of opposite polarity (i.e., positively correlated with the stimulus), consistent with previous findings^82^ were attributed to receptive fields outside the stimulation screen^82^. To exclude these and other noisy ROIs, only those negatively correlated with the stimulus (based on Spearman’s rank correlation) and with ΔF/F₀ values exceeding 0.2 were retained.

#### Full-field flashes analysis

To assess sustained versus transient responses, we calculated the percentage of the peak response retained at the end plateau of the response trace. Peak responses were defined as the difference between the maximum ΔF/F₀ during the stimulus OFF epoch and the mean response 500 ms before stimulus onset. Plateau responses were defined as the difference between the mean ΔF/F₀ during the final 500 ms of the OFF epoch and the same pre-stimulus baseline. The plateau-to-peak ratio was used to characterize response persistence. For the tiny ROIs analysis, multiple small ROIs were manually drawn for each column, and raw fluorescence traces were extracted and provided for each ROI.

#### 11_steps_luminance analysis

To evaluate luminance encoding, plateau responses were calculated as the mean ΔF/F₀ during the first 500 ms of the final second of each stimulus epoch. These responses were fitted with exponential functions, and the decay rates were quantified. Mutual information between luminance and sustained responses was also computed^83^.

#### A_B_steps analysis

Peak responses were defined as the maximum ΔF/F₀ during each B step, relative to the mean baseline activity in the final 2 s of the adaptation period. Sustained plateau responses were calculated as the mean ΔF/F₀ during the last 500 ms of each step. Luminance dependence was assessed by fitting linear regression models to the responses as a function of luminance, and the resulting slopes were used to quantify the sensitivity.

#### Statistics

For all *Drosophila* lamina neuron physiology analyses, means were first computed across ROIs within each fly, and then across flies. Statistical comparisons were performed at the fly level. For normally distributed datasets, two-tailed unpaired Student’s t-tests were used. Normality was assessed using both the Shapiro-Wilk test (*p*>0.05) and the Kolmogorov-Smirnov test (*p*>0.05). For non-normally distributed data, the Mann-Whitney U test (Wilcoxon rank-sum) was applied.

### Mouse tissue preparation

C57BL/6 mice were harvested at postnatal day 18 (P18), and retinas were prepared for analysis as previously described (West et al.^84^). Briefly, retinas were dissected in PBS and fixed for 30 minutes in 4% paraformaldehyde (v/v) with 0.25% Triton-X100 (v/v). After fixation, retinas were washed twice for 5 minutes with 1x PBS, transferred to 7% sucrose in PBS (w/v) for 10 min, and transferred to a 1:1 (v/v) mixture of 30% sucrose in PBS and OCT for 1 hour. Retinas were then frozen in cryomolds and stored at −80 °C.

### Mouse HCR-FISH probe design

cDNA for the target transcript was retrieved from Ensembl^71^ and partitioned into contiguous 52-nt candidate tiles. Each tile was screened for potential off-targets using NCBI BLAST^85^. The 20 tiles with the fewest predicted off-targets were selected as probe templates. For each selected tile, we generated split-initiator probes by bisecting the tile and appending the appropriate HCR initiator sequence (corresponding to the chosen hairpin amplifier) to each half. Oligonucleotides were synthesized by Integrated DNA Technologies (IDT).

### Mouse HCR-FISH

We followed HCR v3.0 protocol from Molecular Instruments^86^ with minor modifications. Following dissection and cryosectioning, tissue sections were rinsed twice in PBS to ensure complete thawing. Samples were incubated in pre-warmed PHB for ≥10 min at 37 °C, which was then replaced with custom-designed probes diluted in PHB (1 μl stock probe solution per 250 μl of buffer) and incubated at 37 °C O/N.

The next day, samples were washed four times for 15 min each with pre-warmed PWB and 5x SSCT with the following ratios: 75% PWB: 25% 5x SSCT, 50:50, 25:75, 0:100. During washes, HCR Amplifiers were heat-activated in a thermal cycler (90 s at 95 °C), and then kept in the dark for 30 min at RT. Samples were washed once with 5x SSCT for 5 min at RT, followed by incubation in AB for ≥30 mins at RT. After cooling, the HCR Amplifiers were diluted in AB (2 ul of 3 uM amplifier stock per 100ul of buffer), added to the samples, and incubated at RT O/N.

The following day, the samples were washed twice for 30 min each in 5x SSCT: first with 100x 405-Wheat Germ Agglutinin, then without. A final wash in 5x SSCT for 5 min was performed before transferring samples to PBS for storage at 4 °C.

## QUANTIFICATION AND STATISTICAL ANALYSIS

The number of L3 neurites per cartridge was evaluated by a permutation test (10,000 permutations). All other image-quantification analyses (except *Drosophila* lamina neuron physiology) were performed in Microsoft Excel and GraphPad Prism version 10.5.0^70^. Group comparisons used unpaired t-test, unless otherwise noted. Data are presented as mean ± SEM. Statistical significance was defined as p < 0.05, with the following notation: \**p*<0.05, \*\**p*<0.01, \*\*\**p*<0.001, ns = not significant. All other relevant statistical information can be found in the figure legends.

*Drosophila* lamina neuron physiology ROI data were averaged per fly before group comparisons. Normality was assessed with Shapiro-Wilk (*p*>0.05) and Kolmogorov-Smirnov (*p*>0.05) tests; normally distributed data were analyzed with unpaired two-tailed t-tests, and non-normally distributed data with the Mann-Whitney U test. \**p*<0.05, \*\**p*<0.01, \*\*\**p*<0.001, ns = not significant.

---

**Figure S1.**
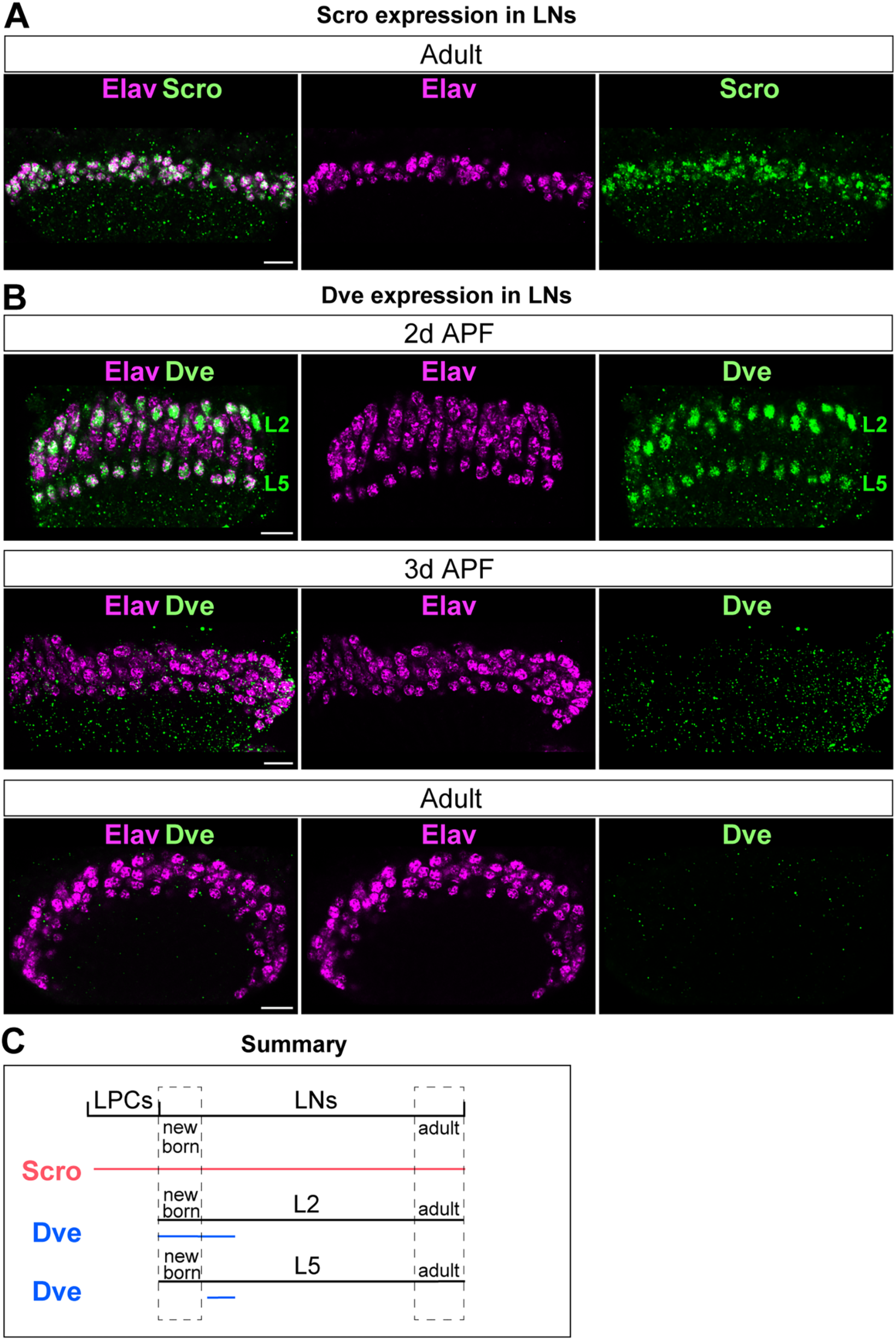
Transient Dve in L2 and L5, and persistent Scro in all lamina neurons, related to Figure 1. (A) Scro persists in all lamina neurons (LNs) in adults. Elav labels LNs. (B) Dve is expressed in L2 and L5 neurons at 2 days after pupal formation (2d APF) but becomes absent by 3d APF and in adults. (C) Schematic summary. Scale bar, 10 µm; n ≥ 5 brains.

**Figure S2.**
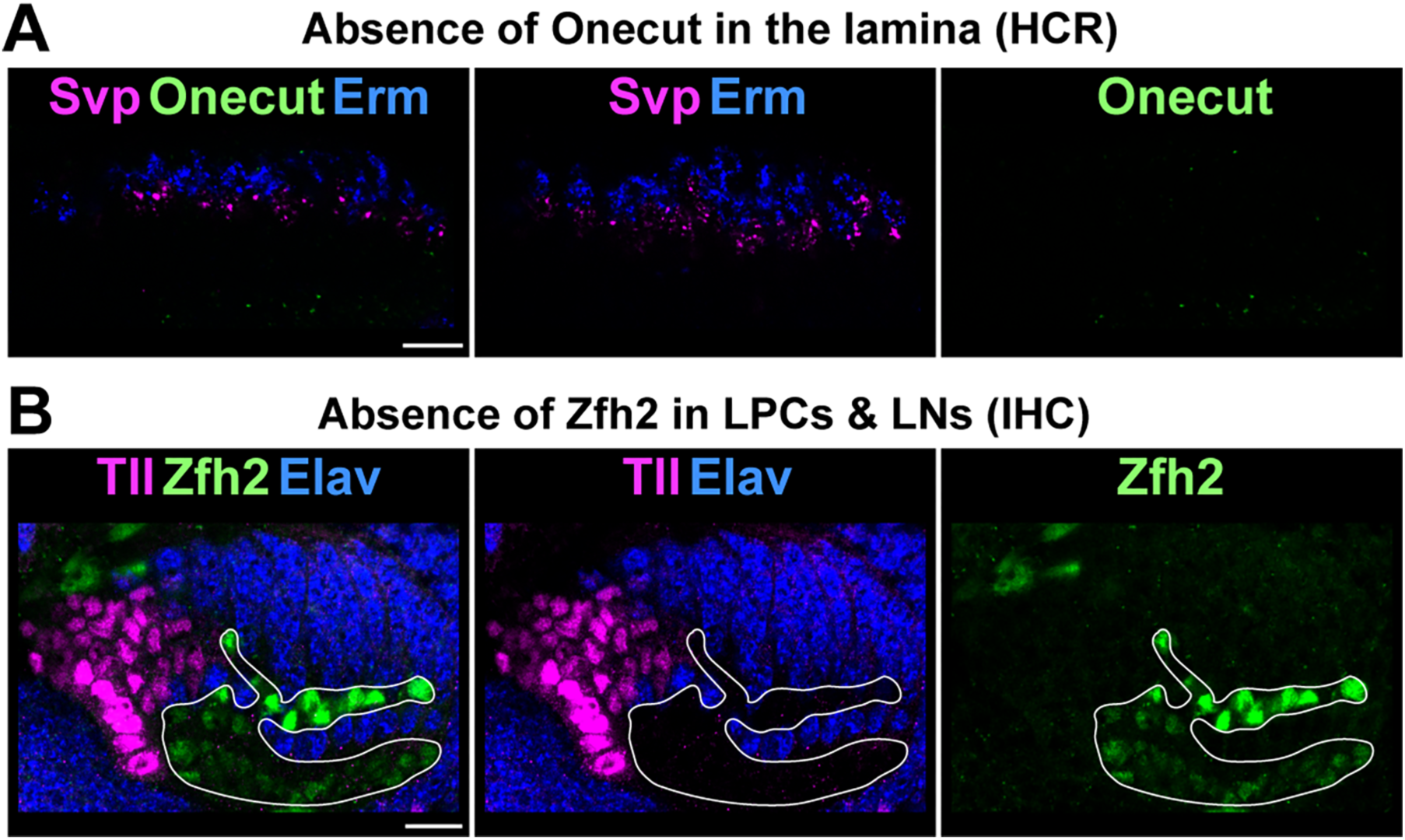
Onecut and Zfh2 are not expressed in the lamina during neurogenesis, related to Figure 1. (A) Onecut mRNA is not detected in the lamina by *in situ* hybridization chain reaction (HCR). Svp and Erm label L1 and L3, respectively. (B) Zfh2 protein is not detected in lamina neurons during lamina neurogenesis. Tll marks LPCs; Elav labels neurons. White outlines: Zfh2^+^ cells. Scale bar, 10 µm; n ≥ 5 brains; sample age: 19h APF.

**Figure S3.**
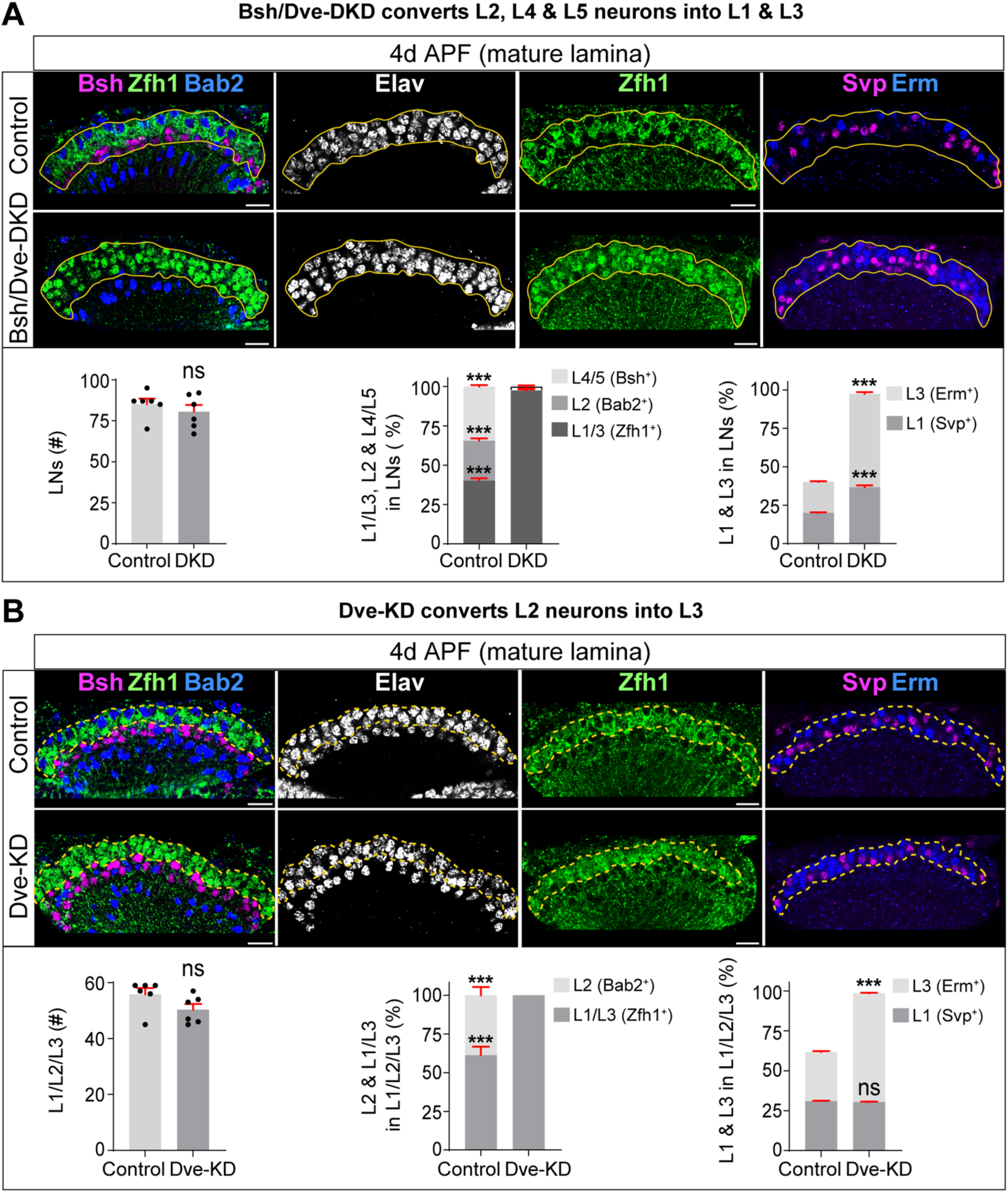
Subtype-specific HDTFs diversify lamina neuron subtypes, related to Figure 3. (A) Bsh/Dve double knockdown (Bsh/Dve-DKD) results in a mature lamina containing only L1 and L3 subtypes. Elav labels neurons; Bab2: L2 marker; Svp: L1 marker; Erm: L3 marker. Yellow outlines: lamina neurons (LNs). Quantification: LN number; percentages of L1/L3 (Zfh1^+^), L2 (Bab2^+^), and L4/L5 (Bsh^+^) within the LN population; percentages of L1 (Svp^+^) and L3 (Erm^+^) within the LN population. (B) Dve knockdown (Dve-KD) results in a loss of L2 neurons and an increase of L3 neurons in the mature lamina. Yellow dashed outlines: L1, L2, and L3 neurons. Quantification: L1/L2/L3 neuron number; percentages of L1/L3 (Zfh1^+^) and L2 (Bab2^+^) within the L1/L2/L3 population; percentages of L1 (Svp^+^) and L3 (Erm^+^) within the L1/L2/L3 population. Data are mean ± SEM; n = 6 brains in (A) and (B); ns, not significant; \*\*\**p*<0.001, unpaired t-test. Scale bar, 10 µm; sample age: 4d APF.

**Figure S4.**
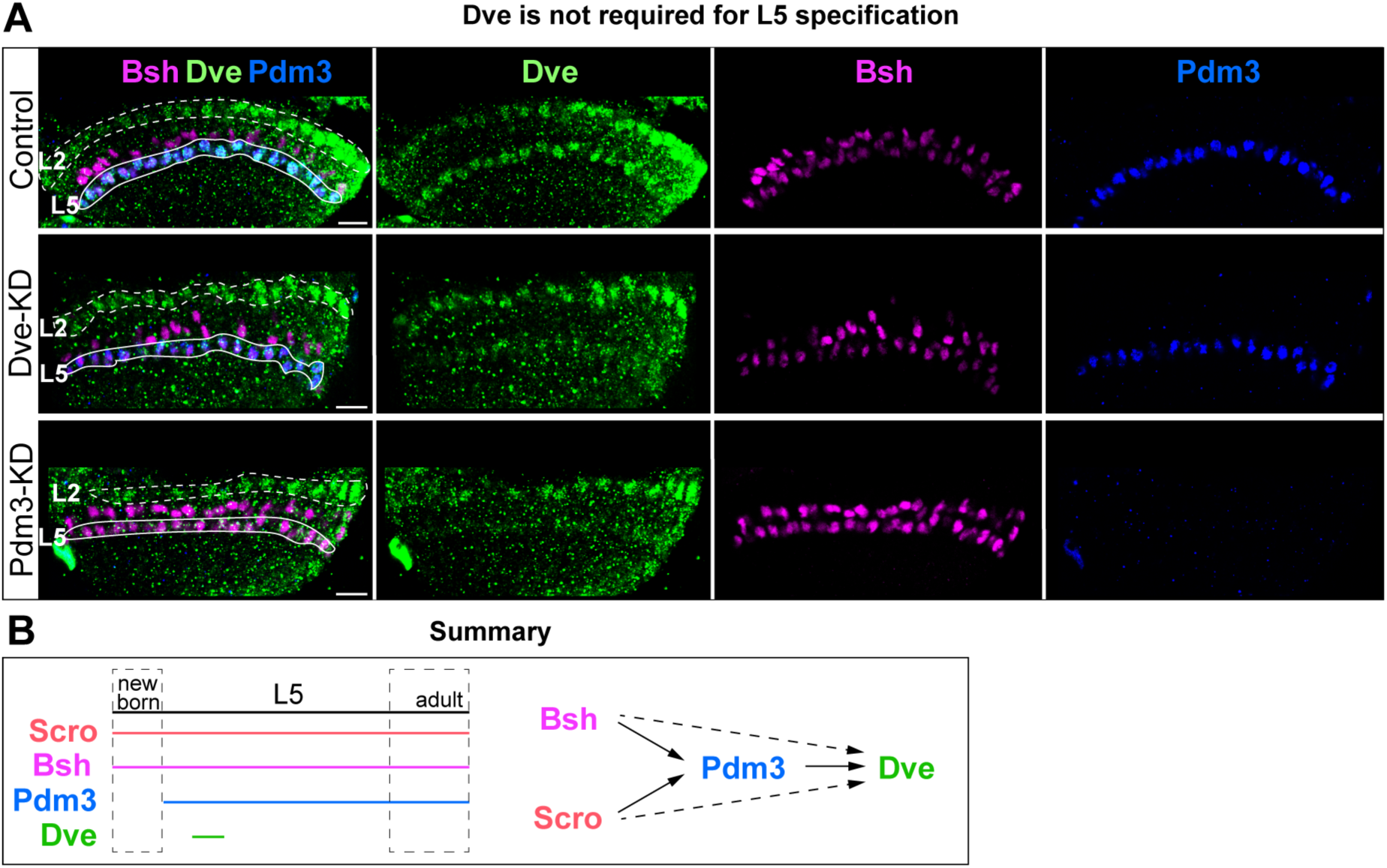
Dve is not required for L5 specification, related to Figure 3. (A) Dve knockdown (Dve-KD) in L5 neurons abolishes Dve expression in L5 neurons but does not affect Bsh or Pdm3 expression. Pdm3 expression becomes absent in Pdm3-knockdown (Pdm3-KD) animals. Pdm3-KD in L5 neurons is achieved with Bsh-L-Gal4 driving UAS-Pdm3-RNAi-#2. White dashed outlines: L2 neurons; White outlines: L5 neurons. (B) Schematic summary. Scale bar, 10 µm; n≥5 brains; sample age: 53h APF.

**Figure S5.**
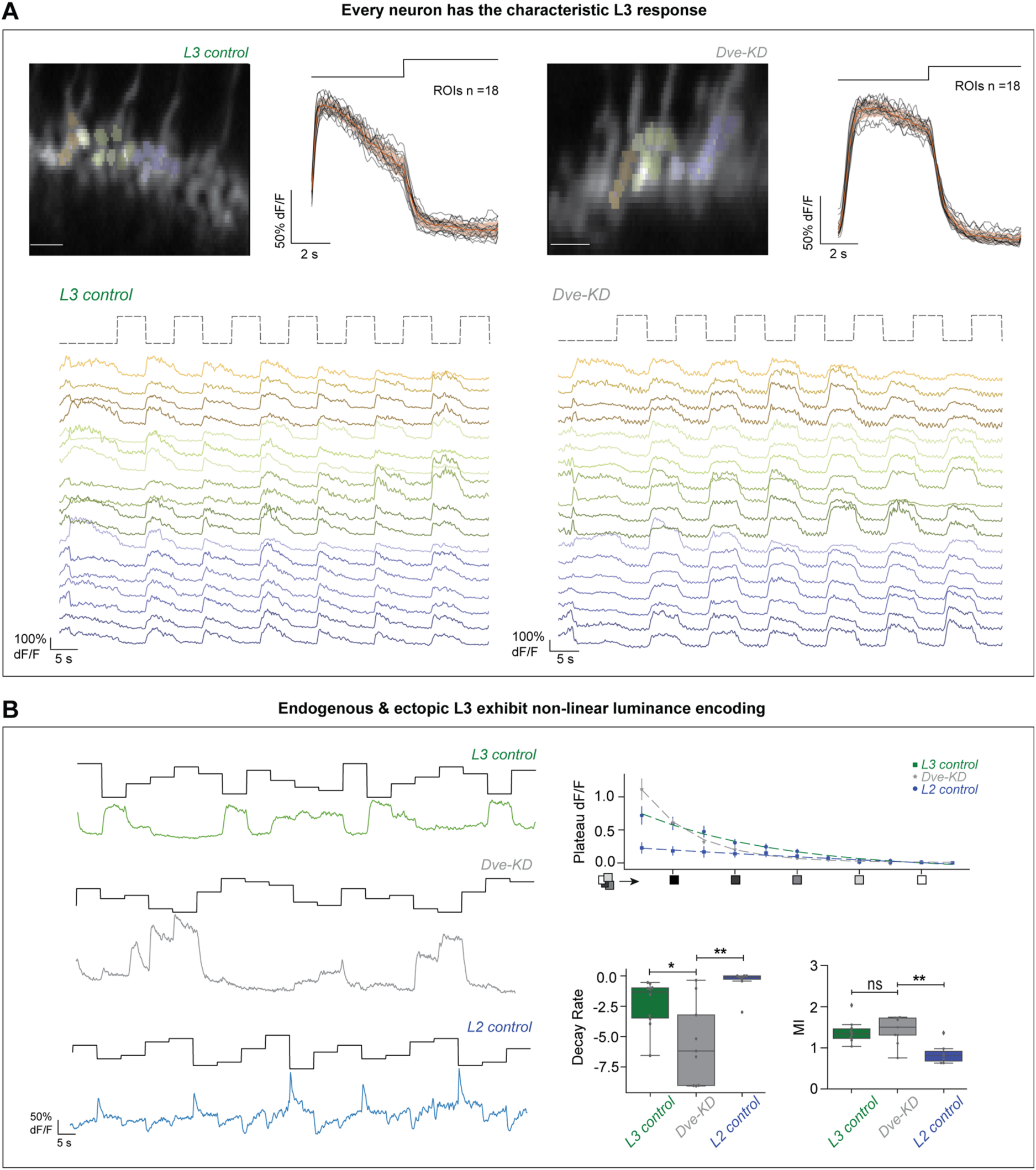
Converted L3 neurons from L2 adopt L3-like sustained response and nonlinear luminance encoding, related to Figure 4. (A) Regions of interest (ROIs) drawn on individual cartridges of L3 axon terminals in L3 control and Dve-KD animals. Average calcium response (orange) and individual ROI responses (black) to full-field flashes in L3 control (n = 18) and Dve-KD (n = 18) animals. Raw calcium traces from individual ROIs in response to full-field flashes in L3 control and Dve-KD animals. Scale-bar, 5 µm; sample age: 5-7 d adult. (B) Representative calcium traces from a single ROI in response to the luminance-steps stimulus for L3 control (green), Dve-KD (grey), and L2 control (blue) animals. Plateau responses plotted as a function of luminance and fitted with exponential functions for L3 control (green), Dve-KD (grey), and L2 control (blue) animals. Quantification of decay rates for the fitted exponential functions: L3 control (n = 10, 82 ROIs; green), Dve-KD (n = 8, 78 ROIs; grey), and L2 control (n = 7, 115 ROIs; blue); \**p*<0.05, \*\**p*<0.01, Mann-Whitney U test. Mutual information between plateau response and luminance: L3 control (n = 10, 82 ROIs; green), Dve-KD (n = 8, 78 ROIs; grey), and L2 control (n = 7, 115 ROIs; blue); ns, not significant, \*\**p*<0.01, unpaired t-test. Sample age: 5-7 d adult.

**Figure S6.**
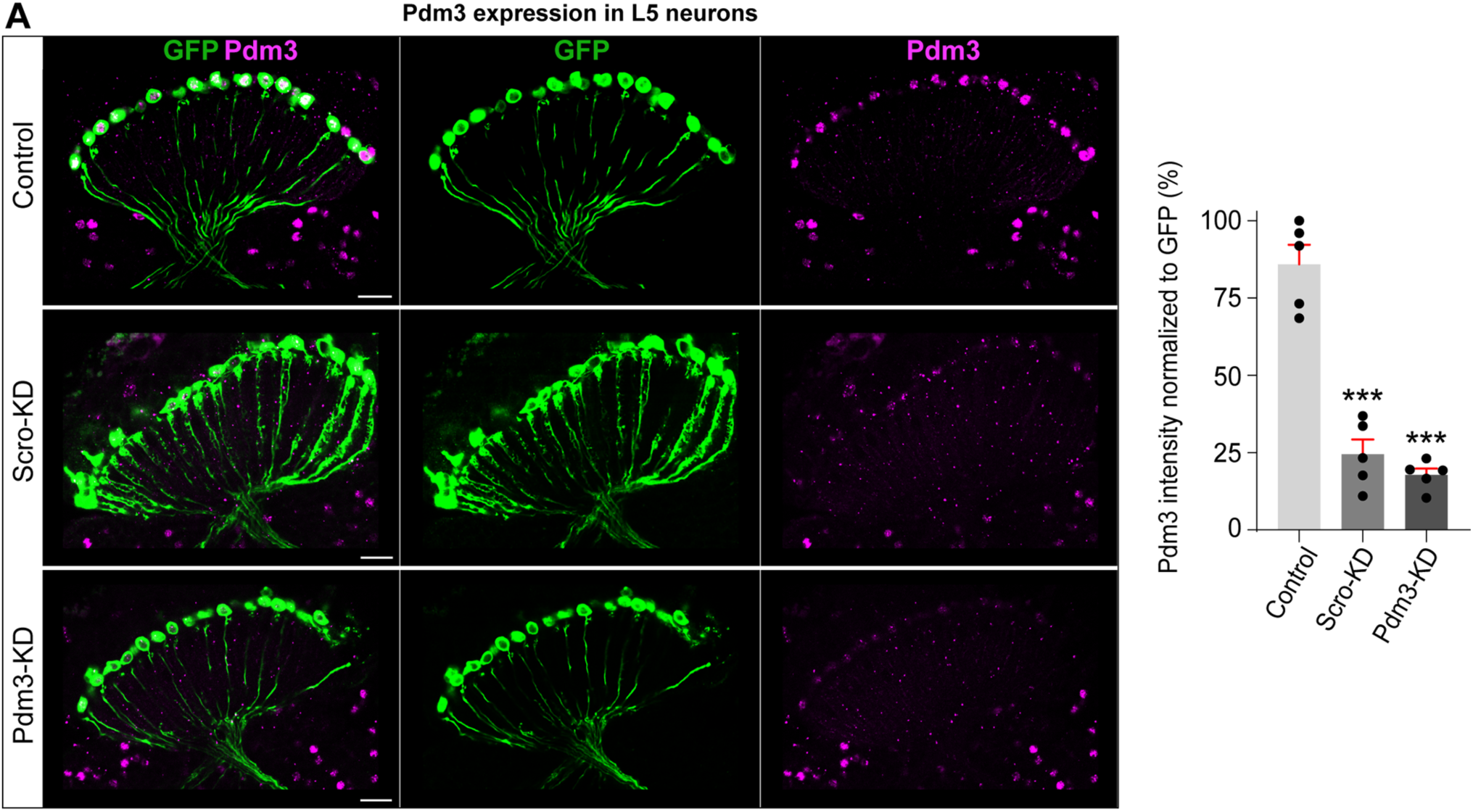
Pdm3 expression is severely reduced in Scro-KD and Pdm3-KD L5 neurons, related to Figure 5. (A) Scro knockdown (Scro-KD) in newborn L5 neurons is achieved with Bsh-L-Gal4 driving UAS-Scro-RNAi. Pdm3 knockdown (Pdm3-KD) in L5 neurons is achieved with Bsh-L-Gal4 driving UAS-Pdm3-RNAi-#1. Bsh-LexA and LexAop-my::GFP are used to trace L5 neurons independent of L5 fate. The Pdm3 signal intensity normalized to GFP is quantified. Data are mean ± SEM; n = 5 brains, \*\*\**p*<0.001, unpaired t-test. Scale bar, 10 µm; sample age: 1d adult.

## Notes

### Competing Interest Statement

The authors have declared no competing interest.

### Summary of Updates

The brain deploys diverse neuronal subtypes to split complex inputs into parallel channels, each tuned to distinct features, enabling rich neural processing. Yet how progenitors generate distinct but functionally related subtypes remains unknown. In the Drosophila lamina (five lamina neuron subtypes receiving photoreceptor input), we uncover the regulatory logic: a pan-class homeodomain transcription factor (HDTF), induced by Hedgehog in progenitors and maintained in all lamina neurons, drives diversification within the lamina neuron class by orchestrating a four-step program across the progenitor-to-newborn neuron transition. Specifically, it establishes progenitor identity, promotes cell-cycle exit, induces subtype-specific HDTFs, and acts as their obligate cofactor to specify distinct subtypes. Loss of subtype-specific HDTFs in newborn, but not older, neurons drives subtype-to-subtype fate conversions at molecular, morphological, and functional levels, including a contrast-to-luminance encoding switch. In the mouse retina, we find that each of the 63 amacrine, 15 bipolar, and 45 retinal ganglion cell subtypes expresses pan-class and subtype-specific HDTFs, indicating evolutionary conservation of this regulatory logic. Given the brain-wide expression of HDTFs across species, these findings convert a longstanding mystery into a testable, generalizable principle for within-class subtype diversification and lay the groundwork for subtype-precise reprogramming and cell replacement strategies.

